# A two-step clockwork mechanism opens a proteo-lipidic pore in PIEZO2 channels

**DOI:** 10.1101/2025.02.10.637531

**Authors:** Shu Li, Tharaka Wijerathne, Aashish Bhatt, Wenjuan Jiang, Jerome Lacroix, Wei Han, Yun Lyna Luo

**Affiliations:** Centre for Artificial Intelligence Driven Drug Discovery, Faculty of Applied Sciences, Macao Polytechnic University, Macao SAR 999078, China; Department of Basic Medical Sciences, Western University of Health Sciences, Pomona, CA 91766, USA; Department of Biotechnology and Pharmaceutical Sciences, Western University of Health Sciences, Pomona, CA 91766, USA; Department of Chemistry, Faculty of Science, Hong Kong Baptist University, Hong Kong SAR 999077, China; Institute of Chemical Biology, Shenzhen Bay Laboratory, Shenzhen 518132, China

## Abstract

Mechanosensitive PIEZO channels are thought to open their pore through tension-induced flattening of large transmembrane arm domains. Yet, the structural basis of this activation remains unclear. Here, we uncover the conformational coupling between arm flattening and pore opening in PIEZO2 by capturing protein motions across length scales using hybrid-resolution molecular dynamics simulations. Sampling multiple microsecond-long trajectories under physiological activation tension show that arm flattening correlates with anticlockwise rotation of the pore domain and with clockwise twisting of inner pore helices, enabling dilation and hydration of a transmembrane pore gate. These clockwork motions enable PIEZO2 to populate two open states with distinct conductance depending on applied membrane tension, in agreement with single-channel electrophysiology. Pore opening is accompanied by the separation of pore helices, creating interhelical gaps which become filled with lipids, resulting in the fully conducting pore being walled by both lipids and protein. The fully open PIEZO2 state recapitulates minimal pore size, conductance, ion selectivity, and outward rectification of chloride currents observed electrophysiologically. This work reveals how tension-induced large-scale rearrangements of the PIEZO2 arms funnel into subtle and dynamic gating motions, providing invaluable structural insights for future structure-function and drug discovery studies.

## Introduction

Mammalian PIEZO channels (PIEZO1 and PIEZO2) play essential roles in a myriad of mechanotransduction processes in vertebrates, including cell volume regulation^1^, neuronal differentiation^2^, epithelium homeostasis^3,4^, development of vascular and lymphatic systems^5–8^, blood pressure regulation^9^, and sensory physiology ^10,11,12,13^. While PIEZO1 channels are present in nearly all types of cells, PIEZO2 channels are mainly expressed in peripheral sensory neurons, in which they encode mechanical stimuli into neuronal signals.

From a structural standpoint, these proteins assemble from three large (∼2500 residues), identical subunits, which delineate a domain-swapped central pore surrounded by three long (∼12 nm) transmembrane domains called arms, or blades. Structures of PIEZO1 and PIEZO2 channels solved by cryo-electron microscopy (cryo-EM) show that these arms naturally curve upward, giving the protein an inverted dome shape^14–18^. This non-planar conformation tends to deforms the lipid bilayer surrounding the channel away from the plane, creating a large PIEZO membrane footprint^19–21^. In these curved structures, a central ion permeation pathway is sealed by hydrophobic residues from three inner pore helices, indicating that the pore is in a non-conducting state.

Several pieces of evidence suggest that the arms serve as flexible, curvature and stretch sensing domains that couple mechanical membrane deformations to pore opening. For instance, atomistic molecular dynamics (MD) stimulations of PIEZO1 show that membrane flattening compel the arms to flatten and the pore to open through side chain rotation of the valine triad^22^. A dilation of the PIEZO1 pore has also been observed computationally through imposing a large membrane tension^23^. Fluorimetric signals from conformational probes inserted along the PIEZO1 arms revealed that motions of the arms and pore opening are temporally correlated^24^. A recent cryo-EM study shows that the PIEZO1 arms fully flatten when the channel is incorporated outside-out into small liposomes due to membrane-protein curvature mismatch^25^. A correlation between osmotically-induced membrane stretch and a physical separation of the PIEZO1 arms was demonstrated using minimal fluorescence photon fluxes microscopy (MINFLUX)^26^, consistent with tension-induced arm flattening. Lastly, the arms of a constitutively active S2472E PIEZO1 mutant adopt a partially flattened conformation in a detergent-solved structure^27^.

In spite of these efforts, it remains unclear how the large-scale PIEZO arms flattening propagate to gate the pore open. Indeed, although PIEZO1 structures in various conformations are available, these structures differ in their spatial resolution in the pore region and lack about one third of the N-terminal arm domain^15–17^. This N-terminal arm region is resolved in the PIEZO2 structure solved in detergent^14^, but no structure or structural model of PIEZO2 in an open or intermediate state exist, making it difficult to piece together a comprehensive gating mechanism for either channel. In this context, this full-length PIEZO2 structure offers a unique opportunity to explore this gating mechanism in details using MD simulations.

Yet, the sheer size of a full-length PIEZO2-membrane system presents a major roadblock for using traditional all-atom (AA) MD simulations to study gating motions at microsecond timescale. To circumvent this timescale issue, previous AA MD simulations of PIEZO1 opening required deploying non-physiological stimuli, such as imposing a very large membrane tension of ∼80 mN m^−1^ which ruptured the membrane within 50 ns^23^, or forcing the membrane footprint of adjacent channels to overlap using periodic boundary conditions (PBC)^22^. Since PIEZO2 requires a higher membrane tension than PIEZO1 to open^28^, opening this channel *in silico* using AA MD simulations under physiological tension is difficult.

Here, we sovled this caveat using an hybrid force field called Protein with Atomistic details in Coarse-grained Environment (PACE). PACE enables simulations of atomistic proteins in coarse-grained (CG) Martini solvent and membrane^29,30^. Unlike Martini protein models, the PACE protein model does not require elastic network restraints to maintain secondary and tertiary protein structure because key anisotropic interactions, such as hydrogen bonding and π-π stacking, are retained between amino acids. In addition, PACE reproduces a tension-induced PIEZO1 open state nearly-identical to the one previously generated using atomistic simulations^22,31^.

Analysis of 20 independent trajectories allowed us to capture a PIEZO2 opening pathway comprising an intermediate subconducting state and a fully conducting state. The relative distribution of subconducting and fully conducting states was tension-dependent in both MD simulations and electrophysiological single-channel recordings. Our simulations reveal a quasi-linear correlation between a large-scale PIEZO2 dome area expansion of ∼600 nm^2^ and a small-scale pore radius expansion of ∼0.2 nm. We further validate the pore size electrophysiologically by measuring the relative permeability of organic cations of different sizes. Importantly, PIEZO2 pore expansion occurs through a series of intricate clockwork-like rotating, tilting, and twisting motions of three outer and three inner pore helices, enabled by tension-induced displacement of the proximal region of the arm. These orchestrated motions cause the hydrophobic pore to expand and become hydrophilic. Large gaps between inner helices created during pore expansion are rapidly filled with lipids headgroups without obstructing the pore, suggesting that the fully conducting pore is walled by both lipids and protein.

## Results

### Membrane size and tension

Using our newly optimized PACE force field^31^, we carried out 20 independent MD simulations (totaling 20.4 µs) of the full-length PIEZO2 channel in a 1-palmityol-2-oleoyl-sn-glycero-3-phosphocholine (POPC) membrane patch of ∼3900 nm^2^ under various constant tensions (**Fig. 1a** and **Movie 1**). This unusually large membrane, relative to the protein size, is important for at least two reasons (discussed extensively in our PIEZO1 study^22^). First, a large membrane is necessary to avoid the PBC effect that flattens the membrane at the boundary and thus destabilizes the curved closed state of PIEZO channels. Second, membrane tension is expected to be laterally isotropic. Yet, PIEZO channels have a C3 symmetry while the membrane patch has a C4 symmetry. This symmetry mismatch could result in a larger membrane bending and expansion force on the PIEZO arms that are closer to the boundary, leading to their asymmetric expansion. In our simulation systems, the minimum distance between the PIEZO arms and membrane boundary is 16.7 nm, which should mitigate this artefact (**Fig. 1a**). Notice that an AA model of PIEZO2 in a solvated 60×60 nm^2^ membrane contains >10 million particles, whereas using PACE, the same system only contains ∼1 million particles without sacrifizing atomistic protein resolution, thanks to the coarse-grained lipids and water model (**Fig. S1a**).

**Figure 1.**
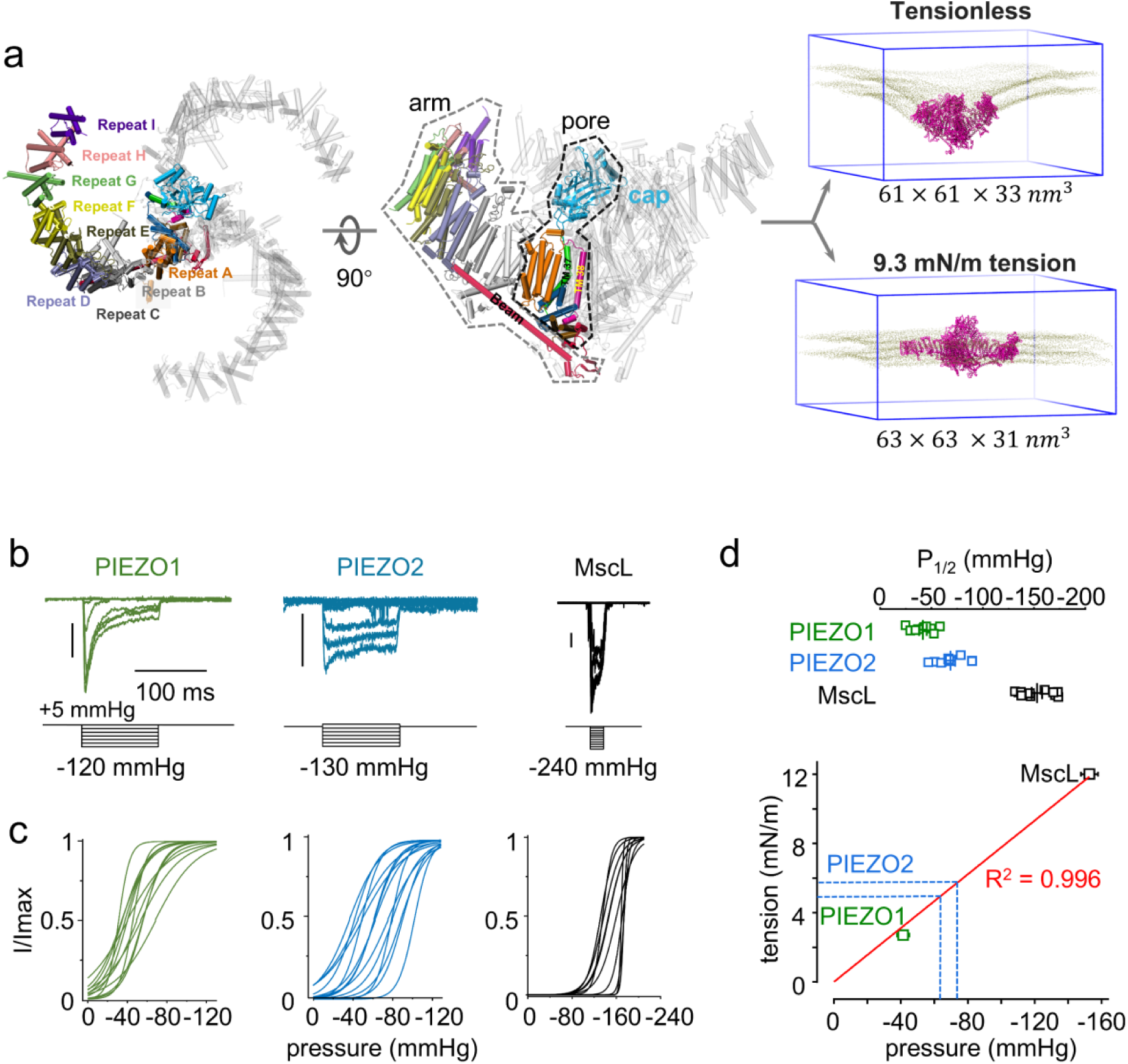
Simulated PIEZO2 system and tension threshold. **a** *Left:* Representation of the PIEZO2 structure showing individual repeats (A to I) in one arm. *Right:* Final conformations of PIEZO2 (magenta) and membrane bilayer headgroup (tan) simulated at 0 and 9.3 mN m^−1^ membrane tension. **b** Representative traces of stretch-activated PIEZO1 (left, green) and PIEZO2 (right, blue) currents at −90 mV. Pipette pressure protocols are shown below each trace. Vertical bars represent 100 pA. **c** Normalized stretch activation of PIEZO1 (green) and PIEZO2 (cyan) represented by Boltzmann-idealized fitted curves of experimental data (not shown for clarity). **d** *Top:* Scatter plot of the half-maximal activation pressures determined for PIEZO1 (green, n=9), PIEZO2 (blue, n=10) and Mscl (black, n=11). *Bottom:* linear fit (red line) between half-activation tension and half-activation pressure with s.e.m. for PIEZO1 (green square) and MscL (black square). Dotted blue lines represent experimental mean ± s.e.m of PIEZO2’s half-activation pressure propagated from the pressure scale to the extrapolated tension scale.

Patch-clamp recordings in which the patched membrane is stretched by application of negative pressure pulses (suction) through the patch pipette show that PIEZO2 is less tension-sensitive than PIEZO1 and produces macroscopic ionic currents with reduced inactivation^28,32^. In our hands, macroscopic currents recorded using the same technique from HEK293T^PIEZO1KO^ cells^33^ transfected with a mouse PIEZO2 plasmid show the same behavior (**Fig. 1b-c**). We estimated PIEZO2’s half-activation tension (tension required to produce 50% of its maximal peak current) by measuring stretch-induced currents from HEK293T^PIEZO1KO^ cells expressing PIEZO1 (whose half-activation tension ranges from ∼1.4 to ∼4.5 mN m^−1^)^34,35^ and MscL (whose half-activation tension is ∼12 mN m^−1^)^36,37^. A linear extrapolation of half-activation pressures measured for PIEZO1 and MscL shows that our membrane patches obey a quasi-linear pressure-tension regime (R^2^ = 0.996) and suggests that stretch-evoked PIEZO2 currents exhibit a half-activation tension ranging from 5 to 6 mN m^−1^ and nearly saturate at ∼9.5 mN m^−1^ (**Fig. 1d**).

In our simulations, we applied a lateral pressure of +1, −2 or −5 bar, which respectively correspond to 0, 9.3 ± 0.2, and 18.0 ± 0.2 mN m^−1^. Our intermediate tension of −2 bar thus corresponds to a physiological near-saturation tension for PIEZO2. We analyzed conformational changes with statistics computed from 4 replicas at 0 mN m^−1^, 9 replicas at 9.3 mN m^−1^, and 7 replicas at 18.0 mN m^−1^ (∼1 microscond per replica). All simulations reached equilibrium, as the root mean square deviation (RMSD) of protein Cα atoms achieved stable plateaux at multiple structural levels, from the global conformation to specific elements such as the inner pore helix (TM38) (**Fig.S1b**).

### PIEZO2 dome under tension

We built our PIEZO2 model based on the full-length PIEZO2 structure (PDB ID: 6kg7) solved in detergent^14^. In this structure, the transmembrane regions of the three arms curve into an inverted dome shape, with an arm angle (averaged angle between the arms and the internal PIEZO axis) of 52 degrees (Fig. 2a). Using the center of geometry (COG) of three inner pore helix (TM38) as reference point, the dome height (measured by the COG of the outer most repeat I) is 8.6 nm, and the height of the extracellular cap domain is 6.0 nm (Fig. 2b). At this conformation, the projected area of the PIEZO2 dome (A_proj_, delineated by the tip of the three arms) is 383 nm^2^ (Fig. 2c). When this PIEZO2 model is embedded in a tensionless Martini POPC bilayer with a bending rigidity of 25.2 ± 2.2 *k*_*b*_T ^38^, the dome height and cap height rapidly decrease and stabilize to 5.5 ± 0.2 nm and 5.8 ± 0.1 nm, respectively. The arm angle increases to 67.8° ± 1.0°, and the projected area expands to 590 ± 51 nm^2^.

**Figure 2.**
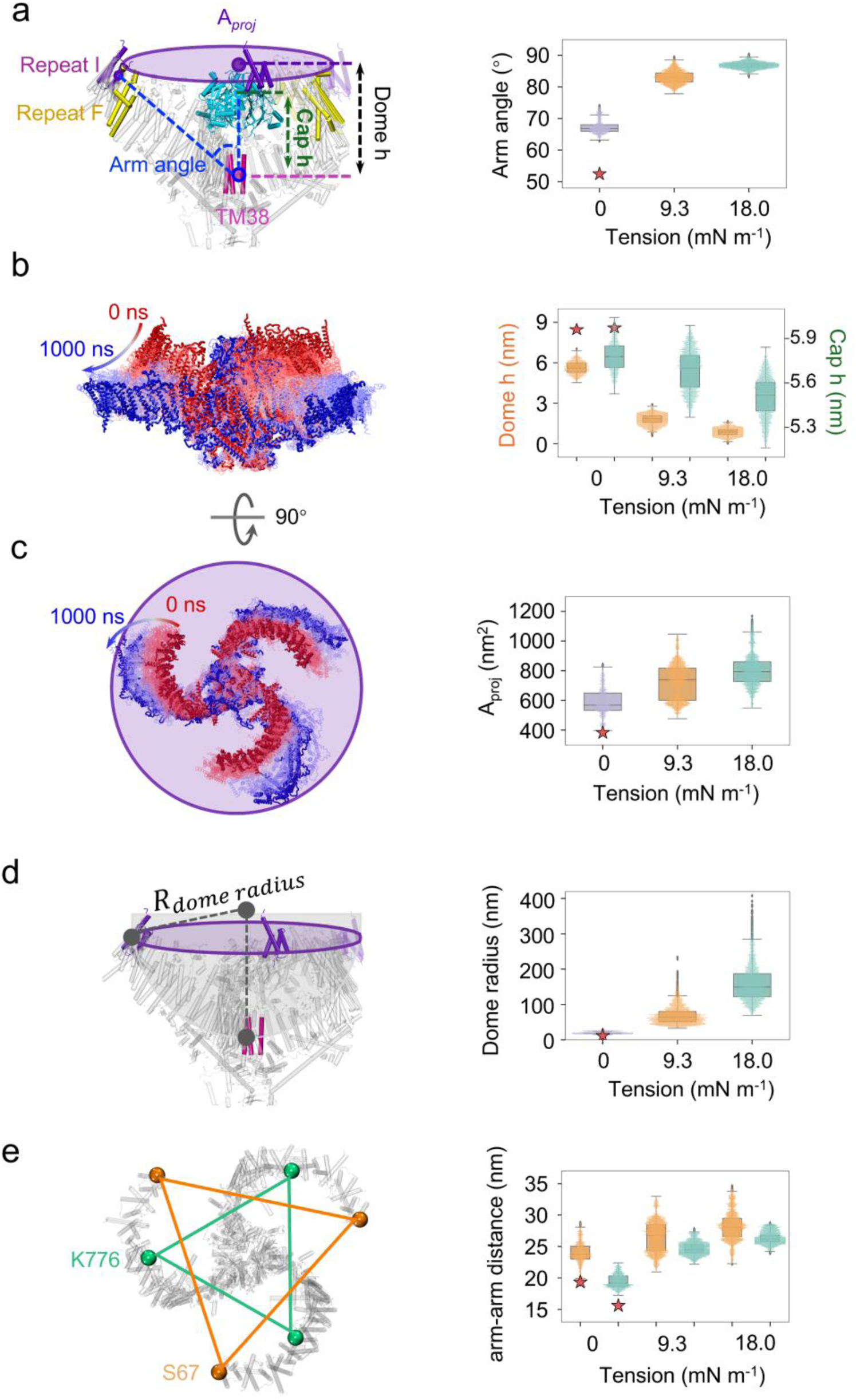
Global tension-induced conformational changes in PIEZO2. **a** The arm angle is defined as the average angle between the arm axis, determined by the COG of the last repeat (repeat I) and inner pore helix (TM38), and the PIEZO2 internal axis, determined by the COG of inner three pore helices and the cap region. **b** Dome height (ℎ) is the COG distance along the Z-axis between three repeat I and three TM38 helices. Cap height is the COG distance along the Z-axis between cap and three TM38 helices. **C** Dome projected area is 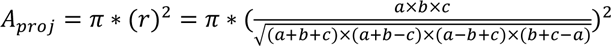, where *r* is the radius of the projected area, *a*, *b*, and *c* are the three distances between the COG of repeat I. **d** PIEZO2 dome radius of curvature 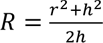. **e** Plots of inter-arm distances between residues S67 (orange) and K775 (green), which correspond to PIEZO1 residues Q103 and D670. In each box plot, a horizontal line indicates the median, while the interquartile range (IQR) spans from the first quartile (Q1) to the third quartile (Q3). Whiskers extend to the smallest and largest data points within 1.5 × IQR, and points beyond this range are considered outliers. Mean ± s.e.m is reported for all values, calculated from 4 replicas at 0 mN m^−1^ tension, 9 replicas at 9.3 mN m^−1^ tension, and 7 replicas at 18.0 mN m^−1^ tension simulations. Conformations sampled during the first 500 ns of each MD trajectory were excluded from our statistical analysis. In each, plot red stars represent corresponding values obtained for the PIEZO2 cryo-EM structure solved in detergent micelles (PDBID: 6kg7).

When a membrane tension of 9.3 mN m^−1^ or 18.0 mN m^−1^ is applied to the system for 1000 ns, the PIEZO2 arms flatten and extend, increasing the arm angle to 83.0° ± 0.5° and 86.8° ± 0.2°, respectively (Fig. 2a). Consequently, the PIEZO2 dome height decreases to 1.8 ± 0.1 nm and 0.9 ± 0.1 nm, cap height decreases to 5.7 ± 0.1 nm and 5.5 ± 0.1 nm (Fig. 2b), and the projected dome area increases, reaching 725 ± 37 nm^2^ and 803 ± 33 nm^2^, respectively (Fig. 2c). By approximating the PIEZO2 dome as a spherical cap, the average curvature radius is 20 ± 2 nm, 69 ± 6 nm, and 160 ± 11 nm at 0, 9.3 and 18.0 mN m^−1^ tension, respecitvely (Fig. 2d).

A recent study employing minimal fluorescence photon fluxes microscopy (MINFLUX) measured the expansion of the PIEZO1 *in cellulo*. Given the structural similarity between PIEZO1 and PIEZO2, these experimental data enable a direct comparison with the simulated structural fluctuation from our MD simulations on a global scale^26^. To this aim, we determined specific PIEZO2 inter-arm distances at various tensions (Fig. 2e). When measured with MINFLUX, the outmost inter-arm distance between PIEZO1 Q103 residues in repeat I is 17.1 ± 4.0 nm in detergent, 25.7 ± 8.6 nm in a resting cell membrane, and 34.7 ± 8.8 nm in osmotically swelled cells^26^. Our simulations show that the inter-arm distances between corresponding PIEZO2 residues S67 follows a similar trend, increasing from 23.8 ± 1.1 nm in tensionless bilayer simulations to 26.6 ± 0.7 nm and 28.2 ± 0.7 nm at 9.3 and 18.0 mN m^−1^. The middle arm distance between PIEZO1 D670 residues on repeat F is 17.8 ± 4.2 nm in a resting cell membrane. The corresponding distances between K775 residues in PIEZO2 was 19.6 ± 0.5 nm in tensionless bilayer simulations, and increased to 24.6 ± 0.3 nm and 26.3 ± 0.3 nm at 9.3 and 18.0 mN m^−1^. The standard deviations of PIEZO2 arm expansion indicate that the middle portion of the arm is more rigid than the outermost arm region (Repeat I H G), consistent with the intrinsic flexibility captured by room-mean-squared fluctuation (**Fig. S2**) and the MINFLUX study^26^. Overall, the triangulated distances measured at two PIEZO2 arm positions in our MD simulations lie well within the standard deviations of the MINFLUX measurements obtained in resting and swelled cells, providing an important validation of global motions captured in our MD simulations.

### Coupling between PIEZO2 dome shape and pore size

The structure of PIEZO2 (PDBID: 6kg7) solved in detergent shows that the pore is constricted by L2743 and F2754 above and below V2750. When simulated in a tensionless membrane, we observed that the flexible side chains of L2743 and F2754 quickly move away from the pore lumen, while all three side chains of V2750 keep pointing toward the central pore axis, forming a stable hydrophobic constriction acting as a main membrane pore gate. Interestingly, V2750 in PIEZO2 corresponds to V2476 in PIEZO1, which also acts as a hydrophobic pore gate that remains closed in both atomistic and PACE simualtions in absence of membrane tension^22,31^. This suggests that both PIEZO homologs use a conserved valine as transmembrane pore gate. The pore size, measured here by averaging the Cβ-Cβ distance between V2750 in all three subunits, was 0.72 ± 0.11 nm in tensionless bilayer simulation, which corresponds to a closed pore. Under both tension conditions, the pore lumen dilates over time, populating two stable sizes of ∼1.3 nm and ∼1.8 nm. Both pore sizes co-exist at different tensions, but they are unequally populated: at 9.3 mN m^−1^, the smaller size is dominant, but at 18.0 mN m^−1^, the larger one predominates (Fig. 3a).

**Figure 3.**
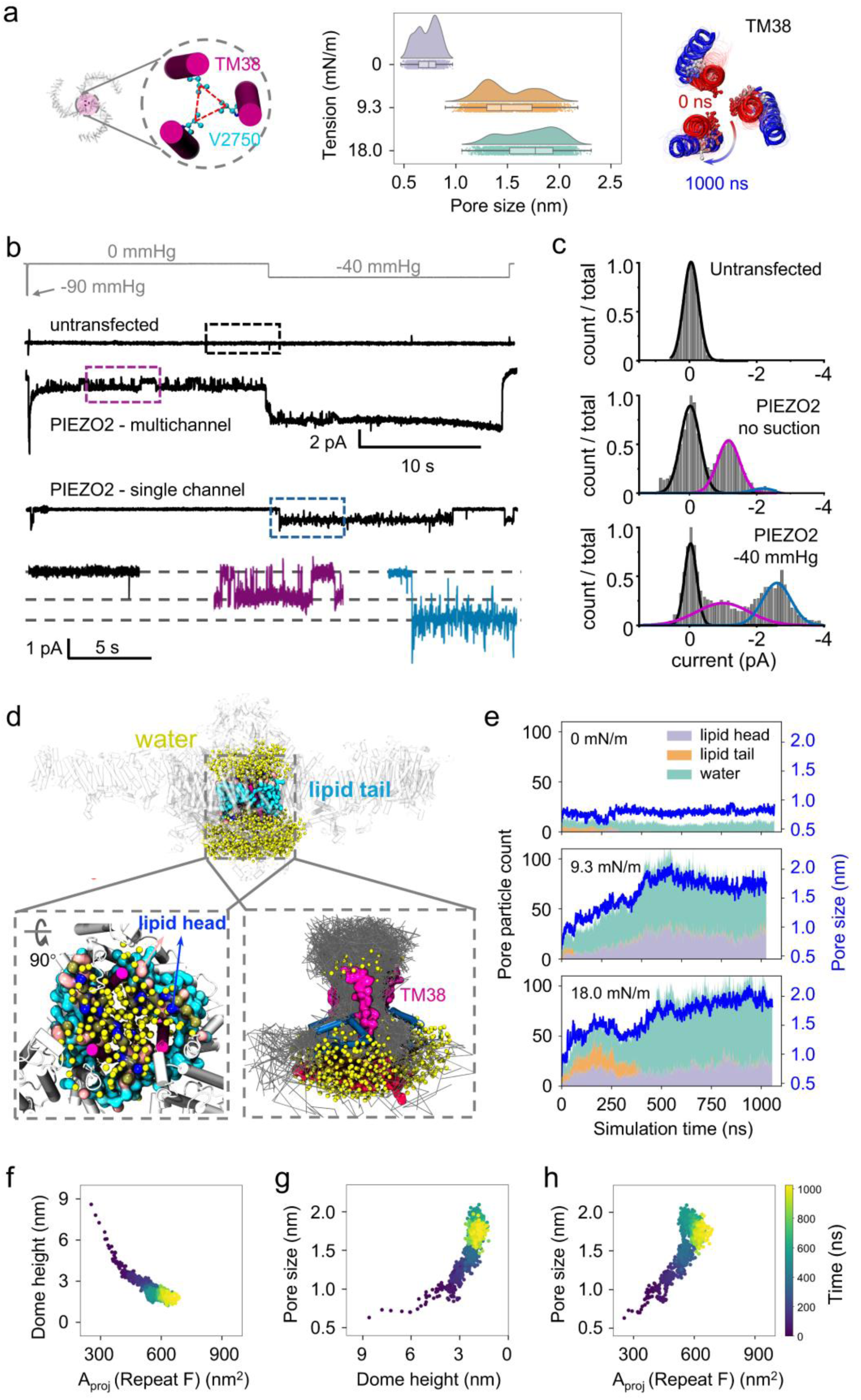
Coupling between dome shape and pore size. **a** PIEZO2 pore size measured by the C*β*-C*β* distance between V2750 sidechains. For simulations at 0 mN m^−1^, error bars represent s.e.m. calculated from 4 replicas. For simulations under tensions of 9.3 mN m^−1^ and 18.0 mN m^−1^, the peak values were obtained using kernel density estimation (KDE). Our statistical analysis only include conformations sampled after 500 ns in each MD trajectory. The right panel shows the overlap of one PIEZO2 trajectory from 0 ns (red) and 1000 ns (blue) under 9.3 mN m^−1^ with V2750 shown in Corey-Pauling-Koltun model. **b** *Top:* Exemplar pressure-clamp traces from untransfected or PIEZO2 transfected HEK293T^PIEZO1KO^ cells. *Bottom:* zoomed view of each insert shown in the above traces. **c** All-point histograms aggregated from four independent recordings and fitted using one (top panel) or three (middle and bottom panels) Gaussian functions. **d** PIEZO2 open state under 9.3 mN m^−1^ tension showing Martini water particles (yellow spheres), lipid tails (cyan), lipid headgroups (blue), TM38 (residues 2740–2761, magenta). The bottom right panel shows pathways of Martini water within the pore (gray lines), the anchor domain (residues 2385–2455, deep blue) and the latch loop (residue 1557–1577, red). See more details in **Fig. S3a**. **e** Pore particle count and pore size at 0, 9.3 and 18.0 mN m^−1^. The number of particles in the pore was calculated within a defined cylindrical region. The radius of the cylinder was determined as the average distance between the *Cα* particles of Val2750 and their centroid, while the height extended 1.5 nm above and below the centroid along the Z-axis (see detail in **Fig. S3a**). Light lavender gray, light orange, and light mint green represent lipid head particles, lipid tail particles, and water particles, respectively. The blue line indicates the pore size (data from 20 replicas in **Fig. S3a**). **f-h** Time evolution of dome height vs. projected area, pore size vs. dome height, and pore size vs. projected area at 9.3 mN m^−1^ (data from 20 replicas in **Fig. S3c-e**).

To experimentally validate theis observation, we performed pressure-clamp recordings from HEK293T^PIEZO1KO^ cells 12-24 hours after transfection with a mouse PIEZO2 plasmid. Patches were held at a constant voltage of −90 mV and subjected to a brief −90 mmHg suction pulse to test if the patch displays activity from one or more channels, followed by 20 s with no suction (0 mmHg) and 20 s with −40 mmHg suction (Fig. 3b). While patches containing only one active channel rarely show discernable opening events at 0 mmHg, they do so at −40 mmHg. This is because PIEZO2 open probability is extremely low at the resting membrane tension of the patch and hence difficult to detect (Fig. 1c). In contrast, multichannel patches show multiple opening events during the 0 mmHg phase of our protocol, as expected because the propability of observing these events increases with the number of channels trapped in the patch. Analysis of these traces and their all-point histograms show that the amplitude of PIEZO2 unitary currents (∼1 pA at 0 mmHg and ∼2.7 at −40 mmHg) yields two distinct conductances (∼11 pS at 0 mmHg vs. 30 pS at −40 mmHg), the larger once being similar to published values for mouse PIEZO2 (∼27 pS^39^ and 23.4 ± 1.14 pS^32^). These results agree well with our simulations and demonstrate that PIEZO2 predominantly populates a subconducting state at the resting tension of the patch (i.e., without external suction), but predominantly populates a fully conducting state in pressurized patches (Fig. 3c).

By analysis how pore dilation correlates with the occupancy of water and lipid molecules in the pore lumen (**Fig. 3d-e and Fig. S3a-b**), we observed that the pore remains closed in tensionless simulations, but dilates in the presence of membrane tension, enabling access to water molecules and to a small number of lipid headgroups, while lipids tails remain confined to the pore wall. For both open states, the pore intracellularly branches out into three lateral fenestrations (Fig. 3d), similar to the multi-fenestrated permeation pathway we and other previously identified in PIEZO1^22,40^.

The time evolution of conformational dynamics from 20 independent trajectories allowed us to decipher the coupling between protein motions across length scales. Under both tested tension conditions, we consistently observed a two-stage dome deformation: membrane tension induces a rapid (<200 ns) PIEZO dome flattening, followed by slow (200-1000 ns) arm extension (**Fig. 3f, Fig. S3c**, and **Movie 1**). Interestingly, during rapid dome flattening, the pore only slightly increases in size and remains nonconducting. But after the dome has nearly flattened, the pore dilates rapidly (**Fig. 3g and Fig. S3d**), increasing with the expansion of the projected dome area (average Pearson’s correlation coefficient from 16 replicas = 0.82 ± 0.02) (**Fig. 3h and Fig. S3e**). This sheds some light on how PIEZO2 enables mechanical force generated from large-scale arm motions to be funneled into subtle rearrangements of inner pore helices.

To further investigate this two-stage gating mechanism, we plotted the time-course of pore dilation against the time course of downward motion for each transmembrane repeat (averaged across the three PIEZO2 arms) for all 20 simulations (**Fig. S3f**). This analysis differs from the global change of the PIEZO2 dome height which we previously inferred from measuring the height of outermost repeat I. Across all 16 tension simulations, the delayed opening of the pore (red lines) tends to correlate with a delayed displacement of the innermost repeat A (orange lines), while all other repeats move in the inward direction in a more uniform manner over time. This suggests that the flattening motion of the PIEZO2 arm can be decomposed into a rigid, monotonic flattening of repeats B-I, followed by a delayed motion of repeat A, which best correlates with pore dilation.

### Clockwork gating motions for PIEZO2 opening

The expansion of the PIEZO2 arms exerts an outward and clockwise (top view) pulling force on the domain-swapped central pore. As a result, the inner pore helices (TM38) rotate counterclockwise relative to the outer pore helices (TM37). We quantified this gating motion by measuring the relative angle between two imaginary triangles: an inner triangle formed by TM38 helices and an outer triangle formed by TM37 helices (**Fig. 4a, Fig. S4a, and Movie 2**). By compiling data across our 16 replicas, this rotation angle decreases almost linearly as the pore dilates from the closed state to the first open (O1) and second open (O2) states, respectively corresponding to the aforementionned smaller and larger open states. The degree of pore hydration and delipidation in O1 and O2 is depicted in the radial distribution function of protein, lipid, and water particles around the pore center axis (**Fig. 4b**). In the closed state, only a small number of water molecules are present above and below the hydrophobic gateformed by V2750. As the pore dilates and rotates into the O1 state, water and lipid headgroups enter the pore, while lipid tails remain outside. Transiting from the subconducting O1 to the fully conducting O2 state, the pore undergoes further dilation and rotation, leading to a significant increase in the number of pore water molecules. In addition to these open states, we also noticed a less populated peak with a similar pore size as O2 but with a larger rotation angle (termed O2’), suggesting that the pore rotation angle is less constrainted at larger pore sizes. Remarkably, the counterclockwise rotation of the inner triangle (**Fig. 4a**) is accompanied by a clockwise twisting of each TM38 helix around their own α-helical axis (**Fig. 4c**). The synchronized twisting of all three TM38 helices rotates six nonpolar sidechains (V2750 and F2754) away from the pore lumen and concomitantly swings in six polar sidechains (S2746 and K2753) into it, enabling the hydrophobic pore to become hydrophilic during its physical expansion, seemingly facilitating delipidation and hydration of the open pore.

**Figure 4.**
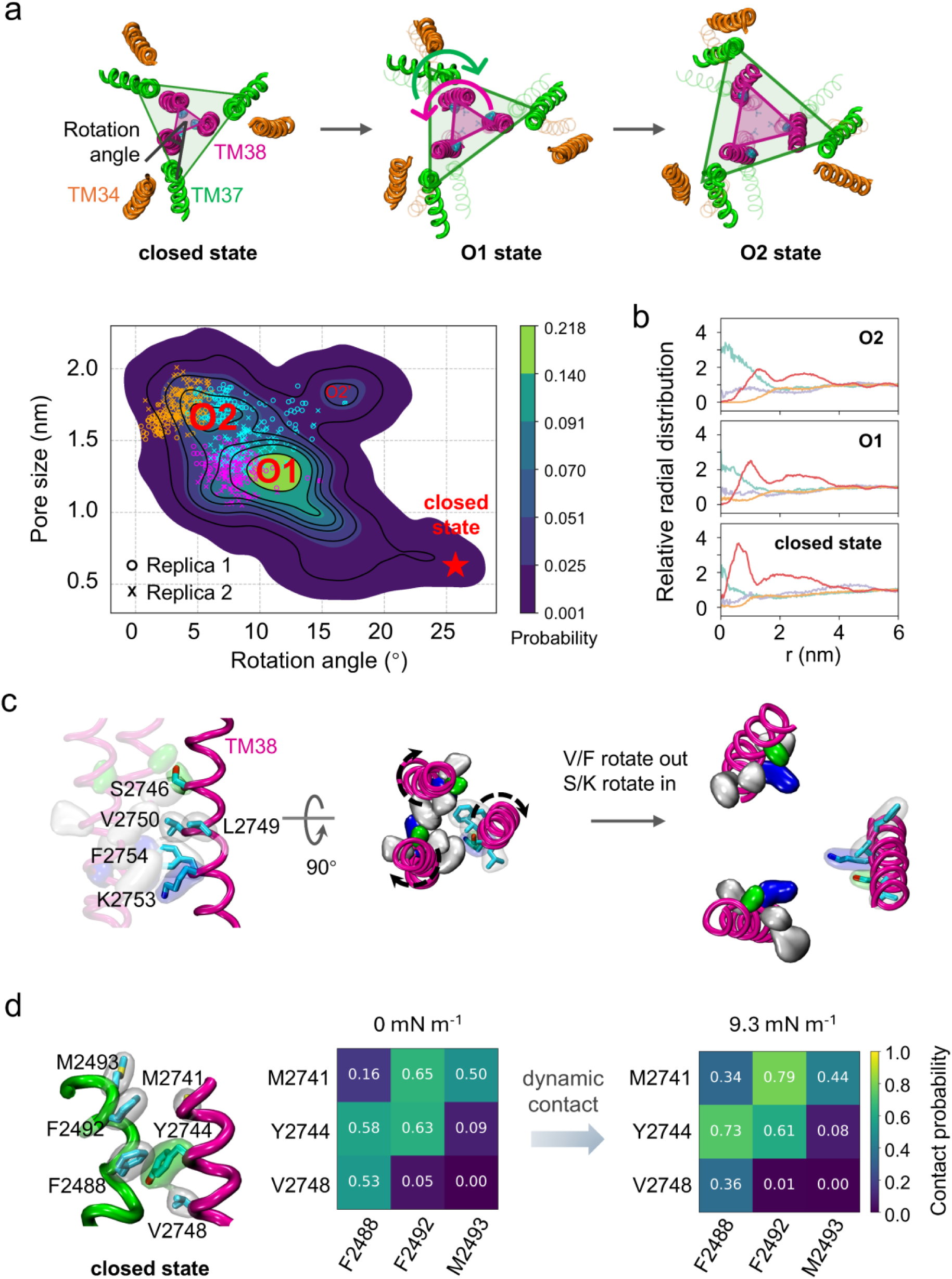
Clockwork PIEZO2 gating motions at 9.3 mN m^−1^. **a** Top view of the pore helix rotation during PIEZO2 opening. The relative angle between the inner and outer triangles, respectively formed by the trios of TM38 and TM37 helices, are plotted against the pore size at 9.3 mN m^−1^ tension using data from all 9 replicas at 9.3 mN m^−1^ tension (**Fig.S4a and Movie 2**). Kernel density estimation was used to calculate the density distribution. Scattered dots in magenta, orange, and blue “o” and “×” symbols represent two 100-ns AA simulations initiated from representative structures sampled from the PACE simulations in the O1, O2, and O2’ states, respectively. **b** Radial distribution function (RDF) of lipid head (light lavender gray), lipid tail (light orange), water (light mint) and protein (red) around pore center. The pore center is defined as the COG of three V2750 Cα particles. The RDF is computed using the pore center as the cylindrical center. The cylinder has a radius of 6 nm and a height of 7 nm (lipid), 5 nm (water), and 3 nm (protein), respectively. For each target particle, the radial distance to the cylinder axis (i.e., the perpendicular distance from the particle to the vertical cylinder axis) is calculated. The RDF, *g*(*r*) is defined as 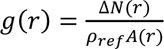, where Δ*N*(*r*): the number of particles within the annular shell between *r* and *r* + Δ*r*; *A*(*r*): the area of the annular shell, calculated as *A*(*r*) = *π*[(*r* + Δ*r*)^2^ − *r*^2^]; *ρ*_*ref*_: the average density of particles in the reference range (4-6 nm). All 4 replicas at 0 mN m^−1^ tension were analyzed for the closed state. All 9 replicas at 9.3 mN m^−1^ surface tension were classified into O1 and O2 states, and RDFs were calculated separately for each state. Only the conformations after 500 ns of each MD trajectory were included in the statistical analysis. **c** Clockwise twisting of TM38 helices rotates nonpolar sidechains out and polar sidechain in the central pore. **d** Hydrophobic contact probability between TM37 and TM38. A contact is defined when the distance between any pair of particles from two residues is less than 0.45 nm. The contact probability was calculated for each simulation from the beginning to the end of the simulation, and the results were averaged across all replicas for each membrane tension. Individual results for all trajectories are provided in **Fig. S4b**.

These clockwork-like gating motions are far from rigid in nature. Indeed, the conformational flexibility of the linker connecting TM37 and TM38 appears critical by enabling these helices to rotate relative to each other. Mechanical force arising from arm flattening is transferred between TM37 and TM38 through dynamic contacts of hydrophobic residues located along TM37 (M2493, F2492, and F2488) and TM38 (M2741, Y2744, and V2748) (**Fig. 4d**). During pore expansion, these hydrophobic sidechains frequently rotate and switch contact partners **(Fig. S4b)**. Disrupting these hydrophobic contacts in our simulations led to the dissociation of TM37 and TM38 during arm expansion. Consequently, force transmission is interrupted and the pore remained closed even after the full arm expansion (data not shown). Therefore, these highly dynamic and “greasy” van der Waals interactions are crucial for enabling TM37 and TM38 to maintain close contact while rotating relative to each other in the clockwork motion.

### PIEZO2 pore size in AA simulations and subconducting state

We further investigated the two PIEZO2 open states captured by PACE under physiological tension (9.3 mN m^−1^) by converting them back to AA models and simulating the pore region containing repeat A, cap, pore, and the C-terminal domain (CTD) using the CHARMM36m force field. We initiated 100 ns AA MD simulations from O1 and O2 states (two replicas each). The conformations sampled atomistically are consistent with the density peaks corresponding to O1 and O2 (**Fig. 4a**, magenta and orange symbols). We also initiated two replicas of 100 ns AA simulation from O2’, a less populated density peak near O2. Both replicas converged back to either the O1 or O2 region (**Fig. 4a**, blue symbols), further arguing that O1 and O2 likely correspond to the main conducting states populated under physiological conditions. Consistent with our PACE simulation results (**Fig. 3d**), the AA simulations show that the gaps between neighboring inner pore helices in O2 state are perfectly sealed by 3 ± 1 POPC lipids (**Fig. 5a**). This arrangement is made possible by the clockwork-like gating motions (**Fig. 4b**) that rotate hydrophilic residues in to promote pore hydration, and concomitantly rotate hydrophobic residues out to seal the pore wall together with lipids.

**Figure 5.**
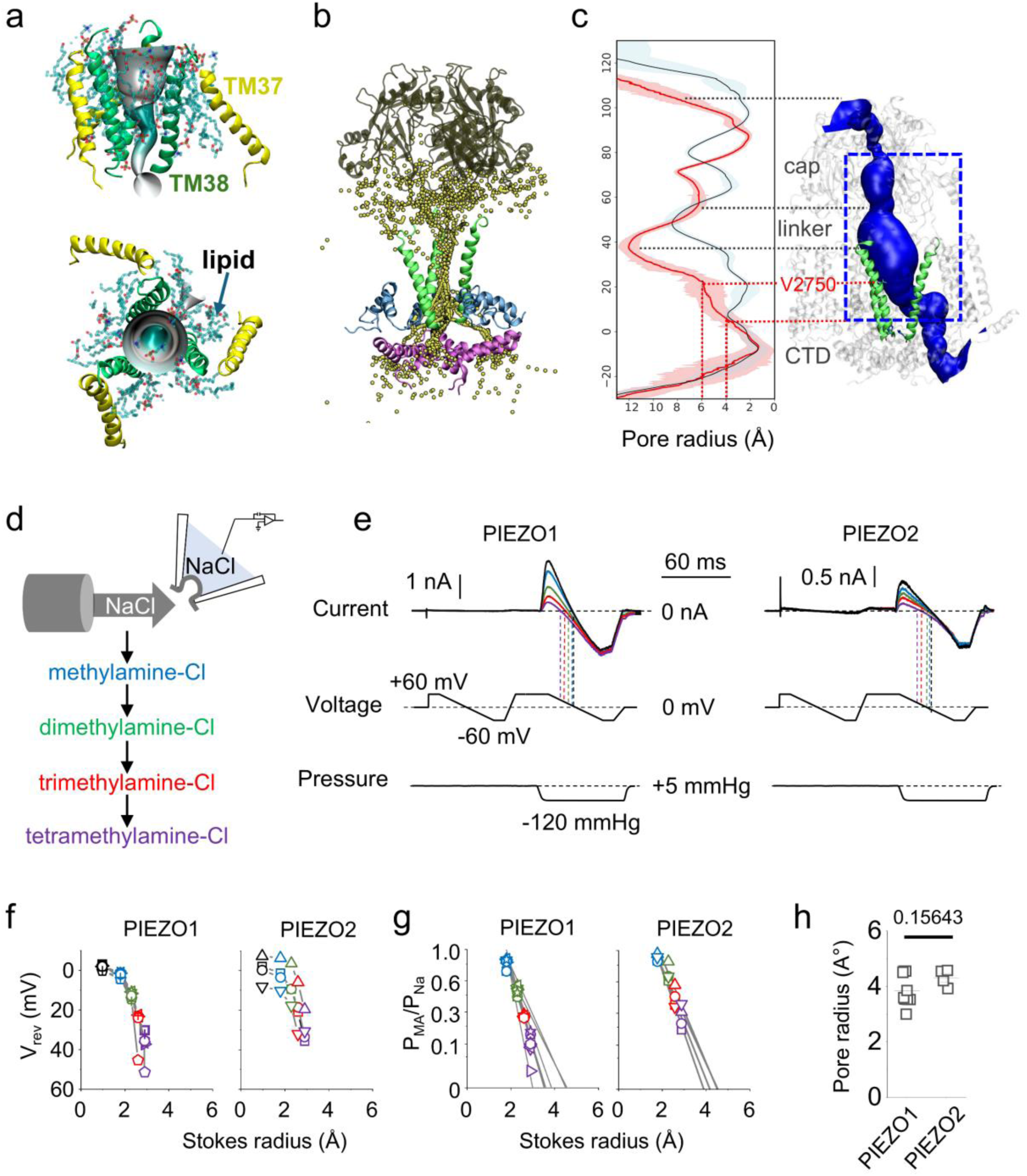
PIEZO2 fully open pore conformation. **a** The backbone of outer pore helix (TM37) is shown in yellow and inner pore helix (TM38) in green cartoon mode. POPC lipids surrounding the pore is shown in transparent licorice mode with atom color code: cyan carbon, red oxygen, blue nitrogen, phosphorus tan. The 3D pore shape is generated using CHAP program. **b** Fenestrated potassium ion pathway illustrated using overlapped potassium coordinates from 100 ns +600 mV AA simulation. **c** Pore radius from 0 mN m^−1^ (blue) and 9.3 mN m^−1^ (red) simulations computed using HOLE program. Standard deviations are shown in shaded color. The corresponding 3D pore profile is shown on the right. The blue square represents the ion accessible path. d Sequence of intracellular saline perfused on to the excised patch. e Exemplar PIEZO1 current (Top) traces generated at indicated voltage (middle) and pressure (bottom) at the presence of sodium (black), methylamine (blue), dimethylamine (green), trimethylamine (red) and tetramethylamine (purple) as the interacellular saline. **f-g** Scatter plots representing reversal potentials (f) and permeabilities for different methylamines relative to Na^+^ as a function of their Stokes radii (data are from singel measurement of 4 independent patches). **h** Scatter plot showing the minimal radius of the open pore calculated for PIEZO1 and PIEZO2 by linear extrapolations (see Methods). The number indicates the exact p-value from a two-tailed unpaired Student’s T-test. Error bars in f-h represent s.e.m.

When applying a constant voltage during AA simulations, we observed a multi-fenestrated ionic conduction pathway, similar to what we observed in PIEZO1^22^. The lower portion of the extracellular cap domain is accessible to water and ions, but the top of the cap remains sealed (**Fig. 5b**). By counting the number of ions crossing the pore, we observed that the O2 state tends to produce a larger conductance than O1 (34.7 ± 2.7 *vs.* 8.9 ± 4.1 pS at −0.6 V; and 38.2 *±* 7.7 *vs.* 11.6 ± 6.2 pS at +0.6 V) (**Table S1**), in good agreement with our experimental results (∼11 pS at 0 mmHg vs. 30 pS at −40 mmHg) (**Fig. 3bc**). The computed O2 state conductance is also similar to published values for mouse PIEZO2 (∼27 pS^39^ and 23.4 ± 1.14 pS^32^). Critically, in the O2 state, more chloride ions contribute to the outward (at +0.6 V) vs. inward (at −0.6 V) current. This result neatly agrees with our recent experimental observation that mammalian PIEZO channels rectify chloride currents in the outward direction when measured in excised patched under stretch conditions, which likely favor PIEZO2’s O2 state (the rectification of chloride currents by the O1 state was not significant)^41^.

The pore radius profile of the O2 state indicates that three pore helices form a funnel shape with a pore radius gradually decreasing from ∼6 Å at the position of V2750, to ∼4 Å immediately above the CTD, i.e. where the permeation pathway turns sideways and split to form lateral fenestrations in each subunit (**Fig. 5c**). To validate these open pore dimensions, we electrophysiologically measured the relative permeability of larger permeants to estimate the minimum pore size for both PIEZO1 and PIEZO2. To this aim, we measured the reversal potential of stretch-evoked macroscopic currents in excised patches, starting from symmetric NaCl solutions to non-symmetric solutions with NaCl in the pipette and organic cations of sequentially larger size in the bath (methylamine, dimethylamine, trimethylamine and tetramethylamine) (**Fig. 5d-e**). The reversal potential values obtained in symmetric and non-symmetric conditions were used to determine the permeabilities of methylated amines relative to sodium ions (corrected for chloride permeation, see Methods), which were plotted against their known Stokes readii. The plots were then fitted with a linear regression based on excluded volume theory to calculate the minimum pore size (see Methods)^42–47^ (**Fig. 5f-h**). The calculated minimum pore radius for PIEZO1 (4.21 ± 0.17 Å) and PIEZO2 (4.35 ± 0.16 Å) were statistically not different (p-value = 0.58925) and match pore size values previously reported for PIEZO1^22^ as well as those reported here for the fully conducting PIEZO2 open pore (∼4 Å, see **Fig. 5c**).

## Discussion

We report here a realistic computational model for PIEZO2 activation by membrane stretch, extensively validated by electrophysiological measurements of tension-sensitivity, conductance, ionic selectivity, and open pore size. To circumvent the inherent challenges associated with computational simulations of large protein-membrane systems^48^, we used PACE, a hybrid force-field that drastically reduce the number of atoms in membrane and solvent while maintaining atomistic details of protein motions. PACE allowed us to consistently capture realistic opening transitions of a full-length PIEZO2 channel embedded into a large membrane patch under a physiologically-relevant membrane tension. In contrast to previous atomistic simulations of PIEZO1^22,23^, PACE obviates the need to minimize the size of the simulated system (which truncates the PIEZO membrane footprint and introduces PBC artefacts), and/or to use a non-physiological membrane tension (which quickly ruptures the lipid bilayer). The large membrane deformation associated with the gating of PIEZO channels could be fully captured by our multiscale PACE simulations. Critically, the enhanced speed of multiscale simulations allowed us to generate many independent replicas, enabling the statistical identification of gating-relevant motions from random thermal fluctuations.

The tension-induced expansion of two triangulated arm-arm distances in our PIEZO2 simulations matches the expansion of homologous arm-arm distances captured in PIEZO1 using MINFLUX in resting and swollen cells^26^. Changes in the PIEZO dome geometry also show similarities between our PIEZO2 simulations and experimental estimations of the PIEZO1 dome. Using membrane elastic theory and cryo-electron tomography, Haselwandter and colleagues reported a intrinsic dome curvature radius of 42 ± 12 nm (and a bending stiffness of 18 ± 12 *k*_*b*_T) for PIEZO1 embedded into a lipid vesicle^20,21^. On the other hand, the Xiao group reported cryo-EM structures of PIEZO1 reconstitued into small liposomes with extreme curvature radii of ∼10 nm (curvature match) and ∼117 nm (curvature mismatch)^25^, and intermediate radii ranging from ∼14 nm to ∼32 nm for a PIEZO1 gain-of-function mutant^49^. In our simulated POPC lipid bilayer (with a bending rigidity of 25.2 ± 2.2 *k*_*b*_T), the PIEZO2 curvature radius increases from 20 ± 2 nm at 0 mN m^−1^, to 69 ± 6 nm at 9.3 mN m^−1^, and to 160 ± 11 nm at 18.0 mN m^−1^ (**Fig. 2d**). Our PIEZO2 dome’s radius of curvature of 20 ± 2 nm in a POPC membrane at zero tension thus lies between PIEZO1 experimental curvature values reported above. By approximating the PIEZO2 dome as a spherical cap, we also computed its total surface area based on dome height and curvature radius. Our atomistic simulations show that the PIEZO2 dome surface increases with tension, from 685 ± 52 nm^2^ at 0 mN m^−1^, to 736 ± 36 nm^2^ at 9.3 mN m^−1^ and to 805 ± 33 nm^2^ at 18.0 mN m^−1^. This is largely due to the intrinsic flexibility of PIEZO’s peripheral arm region, reflected by the high mobility of repeats I H and G (**Fig. S2a, Move 1**), enabling the protein to further unfurl its arms at higher tensions.

The unfurling of the PIEZO2 arms correlates with an outward and clockwise (as seen from the top) rearrangement of the domain-swapped central pore domain. During this motion, the triangle formed by the inner pore helical trio (TM38) rotates counterclockwise. Concomitantly, each TM38 helix also individually undergoes a clockwise twisting motion around their own α-helical axis. These orchestrated motions dilate the pore and change its chemistry from hydrophobic to hydrophilic, leading to a subconducting open state that is predominantly populated at low membrane tension in both electrophysiological recordings and simulations. Further arm expansion correlates with further separation of TM38, enlarging the pore into a fully conducting state.

Concomitantly to pore dilation, three pore helices separate from each other, creating large interhelical gaps granting access to the pore lumen by POPC lipids. These gaps seem to emerge due to PIEZO’s trimeric organization, resulting in a loose packing of three pore helices. Indeed, the homotrimeric acid-sensing ion channel 3 (ASIC3) displays similar gaps when they open, enabling POPC lipids to partially occlude the open pore and reduced single channel conductance^50^. In our PIEZO2 simulations, however, pore occlusion by POPC lipids did not seem to occur, likely because pore hydration predates the formation of these gaps. Pore hydration keep most POPC lipids to the periphery of the pore lumen, suggesting that bulk lipids act as *bona fide* molecular constituents of the open pore wall. A similar proteo-lipidic open pore was recently observed for members of the OSCA/TMEM63 channel family^51^.

The ability of bulk lipids to diffuse in and out of channel pores through membrane fenestrations has been proposed to contribute to tension-dependent gating of certain mechanosensitive ion channels, including OSCA/TMEM63^51^, MscS^52^ and TRAAK^53^. Although previous computational studies have reported that the PIEZO1 pore can be fully occluded by lipids^22,23^, we recently showed that lipid density in simulated channel pores may be overestimated due to an overlooked simulation artefact that decreases pore hydration^54^. Hence, our clockwork gating mechanism seems to rule out a mechanism in which PIEZO2 primarily senses membrane tension through the ability of bulk lipids to move in and out of the pore lumen. Yet, we do not rule out the possibility that lipids other than POPC may alter channel function by occluding the pore. Indeed, PIEZO1 and/or PIEZO2 channels are robustly modulated by many lipids, including cholesterol^55–58^, dietary fatty acids^59,60^, phosphatidic acid^61^, phosphoinositides^62,63^, ceramides^64^, and phosphatidylserine^65^. In addition, recent structures of human PIEZO1 complexed with a lipidated regulatory protein shows lipid densities in the pore^66^. Atomistic simulations using our PIEZO1 and PIEZO2 open states in the presence of regulatory lipids should allow us to test whether lipid modulation of PIEZO channels arises from changes in membrane mechanical properties and/or from specific lipid-channel interactions.

Although similar tension-induced gating motions occur in PIEZO1 and PIEZO2, including pore dilation, valine side chain rotation, and dome flattening^22,31^, the chronology of these events vary between the two channels. Indeed, in PIEZO2, the membrane gate remains closed during initial arm flattening, opening subsequently as the arms unfurls outwardly. In contrast, the PIEZO1 pore opens readily upon arm flattening^22,23^. This inherent difference may explain why the tension threshold to open PIEZO2 is higher than PIEZO1^28^.

Modulation of PIEZO2 channel activity with small molecules represents a promising approach to treat a number of ailments, ranging from certain forms of pain associated with inflammation^67–69^ to gastro-intestinal disorders^70^ and incontinence^71^. While no known PIEZO2 selective inhibitor or activator currently exist, we anticipate that our PIEZO2 open state model and gating mechanism will facilitate the identifications of novel chemical modulators through virtual, structure-guided drug discovery pipelines. A similar approach led us to identify an allosteric agonist binding site in PIEZO1, sandwiched between repeats A and B, and two novel PIEZO1 activators^72–74^. Interestingly, we found that this region acts as a hinge during PIEZO2 arm flattening: the bulk of the arm, formed by repeats B-I, flattens rapidly under tension but repeat A movement occurs with a delay, as does pore dilation. Hence, we anticipate that small molecules wedging between PIEZO2 repeats A and B might also modulate PIEZO2 function.

### Limitations of our study

PIEZO2 may use distinct mechanisms to detect different types of mechanical stimuli. For instance, an intrinsically disordered intracellular domain of PIEZO2 is required for its activation by cytoskeleton-transmitted forces, but not for its activation by stretch^32^. Our PIEZO2 simulations without intracellular domains and cytoskeletal components thus only probe the mechanism of stretch activation through opening of a membrane pore gate. Along a similar line, structural studies of PIEZO1 suggest the existence of multiple permeation gates beside the transmembrane pore gate, including a lateral gate located at intracellular fenestrations^25,40,49^. Since structural determinants of these additional gates are unresolved (or absent) in the PIEZO2 structure, we were unable to study their role in stretch activation.

## Methods

### PIEZO2 atomistic model preparation

The PIEZO2 model was derived from the cryo-EM structure of full-length mouse PIEZO2 in closed state (PDB ID: 6kg7). PIEZO2 homotrimer is composed of 114 transmembrane helices, with each monomer contributing 38 transmembrane helices. Among the unresolved parts in PIEOZ2 structure, three extracellular loops (residues 1383-1426 on PIEZO repeat C or THU7, 2056-2074 on repeat B or THU8, and 2283-2298 on repeat A or THU9) exhibit more than 60% sequence similarity to equivalent segments in PIEZO1, whose structures have been resolved in multiple conformations in varying resolutions. Thus, we utilized multiple PIEZO1 structures (PDB ID: 6B3R and 7WLU) as templates for homology modeling of these loops using the MODELLER v10.4 program^75^. The model also lacked linker loops between the cap domain and the outer and inner pore helices (TM37 and TM38) at residues 2493-2500 and 2732-2738. These were constructed using the loop modeling module within MODELLER. Additionally, we modeled 43 residues (1515 - 1557) connecting the C-terminal of the beam to the Latch. The final PIEZO2 model contains 1953 residues (8-79, 122-142, 201-304, 332-358, 491-615, 675-823, 934-1577, 1668-1727, 1948-2111, and 2236-2822) per monomer and a total of 5859 residues for the entire homotrimer. Remaining residues, primarily the cytoplasmic intrinsic disordered regions lacking corresponding template structures, were not modeled. Additionally, the PDBFixer program^76^ was used to repair missing heavy atoms in the PIEZO2 model.

### New approach to construct dome-shaped membrane

In our previous MD simulation studies of the truncated PIEZO1 cryo-EM structure^22,31,73^, we positioned the PIEZO1 protein within an AA or Martini model of flat lipid bilayer. Over extended equilibrium simulations with protein backbone constraint, the membrane naturally developed a dome structure encompassing the PIEZO1. However, for the full-length closed PIEZO2 cryo-EM structure, the transmembrane region of the peripheral arms are above the extracellular cap domain, forming a much deeper dome structure, one that could not be fully encapsulated by the spontaneous membrane curving through long MD simulations. To tackle this problem, we developed an approach to efficiently construct a membrane topology conforming to the unique of the protein (**Fig. 6**).

**Figure 6:**
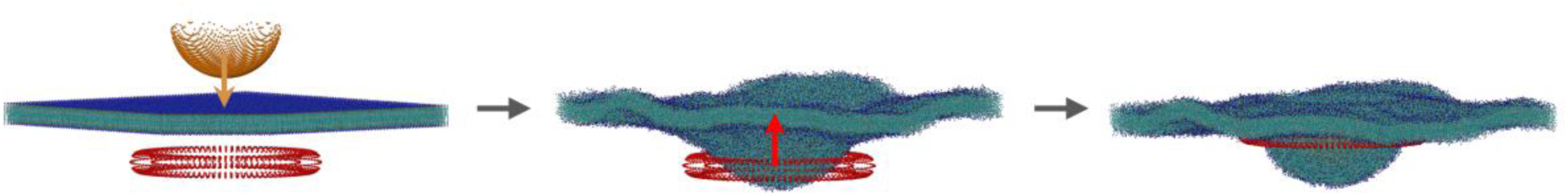
Approach to construct dome-shaped membrane. 1. First, based on the geometric shape of the PIEZO2 dome, we created a hemisphere structure comprising 2129 dummy particles (dome shape), and a torus structure with a minor radius of 2 nm and a major radius of 16 nm, made up of 1800 particles (doughnut shape). These two structures were positioned 16 nm apart along the Z-axis. 2. Next, we constructed a 60 nm × 60 nm × 44 nm flat Martini2 POPC bilayer membrane using the insane.py script. The hemisphere and torus were then integrated into the POPC bilayer system. The hemisphere was positioned directly above the lipid layer while the torus was positioned directly below it. We added force field parameters using Lennard-Jones potential with C6 = 0 and C12 = 0.01 to induce repulsive interactions between the dummy particles (including the hemisphere and torus) and the Martini POPC particles. 3. Finally, we kept the position of the torus fixed, designating the Z-axis as the reaction coordinate, and applied a force constant of 100,000 kJ mol^−1^ nm^−2^ to pull the hemisphere downward by 15 nm. Similarly, with the hemisphere fixed, the same force was applied to push the torus upward until the membrane structure resembles the PIEZO2 shape, and removed the structured particles (code available at https://github.com/LynaLuo-Lab/Piezo2).

### PIEZO2 CG Martini system construction and simulation

We converted the atomistic PIEZO2 structure into the standard CG Martini2 elastic network (ELNEDYN22) model^77^ with a 0.9 nm pairwise diatance cutoff and a 500 kJ·mol^−1^·nm^−1^ force constant, and integrated it into the pre-formed dome-shaped Martini POPC membrane. The system was solvated and neutralized using a 150 mM NaCl solution. The final dimension of the PIEZO2 system is 60 nm × 60 nm × 35 nm, containing a total of 1,069,143 Martini beads. The CG Martini simulation was carried out using Gromacs (version 2022)^78,79^. Initial energy minimization was conducted for 5000 steps, followed by pre-equilibrium simulations lasting 4.75 ns., During the initial water relaxtion step, the whole lipids was restrained to avoid exccesive pore lipids according to our published protocol^54^. Afterwards, positional constraints on the protein’s backbone beads were gradually relaxed from 1000 kJ·mol^−1^·nm^−2^ to 500 kJ·mol^−1^·nm^−2^. For nonbonded interactions, a cutoff of 1.1 nm was employed for both short-range electrostatic and van der Waals forces. Long-range electrostatic interactions were calculated using the default reaction-field algorithm. The simulations were performed with a time step of 20 fs. Temperature was maintained at 323 K using the V-rescale thermostat, and pressure was regulated at 1 bar using a semi-isotropic coupling method implemented via the Parrinello-Rahman algorithm. The CG simulation extended over 1 µs, with positional restraints (500 kJ·mol^−1^·nm^−2^) maintained on the PIEZO2 backbone beads to allow for the relaxation of other system components, particularly the lipid molecules. The final Martini conformation was used to construct the PIEZO2 PACE model.

### PIEZO2 PACE system construction and simulation

We used the atomistic PIEZO2 model to construct its united atom (UA) structure and parameters, employing the recent PACE force field specifically designed for membrane protein simulations and benchmarked on PIEZO1 model^31^. We then replace the simulated PIEZO2 Martini model with the PACE model by aligning the protein backbone beads, retaining both the Martini membrane and solvent components from the equilibrated system. The PACE system comprises 5,859 protein residues, 11,355 POPC lipids, 896,114 CG water particles, 11,390 CG Na^+^ particles, and 11,483 CG Cl^−^ particles, totaling 1,110,168 particles, that is only 3.8% increase from Martini system (**Fig. S1a**).

In recent PACE force field optimization, we reported that increasing the transmembrane helical backbone hydrogen bonding strength by 1.5-fold or 2-fold resulted stable membrane protein (overall protein RMSD of 3.24 ± 0.12 Å and 3.30 ± 0.05 Å, repectively) and good agreement in the number of hydrogen bonds in helical regions with AA simulations. Given the structural integrity of helices is crucial for mechanotransduction, the PACE force field with 2-fold hydrogen bonding strength within the membrane helices is chosen for simulating PIEZO2 under mechanical stress.

All PACE simulations were carried out using Gromacs (version 2021 and 2022). The system underwent an initial energy minimization over 5000 steps, followed by a 1.2 ns pre-equilibrium phase which strong positional constraints (500 kJ·mol^−1^·nm^−2^) were applied to the protein’s Cα atoms. Next, 50 ns equilibration simulation was proceeded with positional constraints applied only to the Cα atoms of cryo-EM resolved residues (excluding the homology models of the loops and linkers described above). Afterwards, weak positional restraints of 50 kJ·mol^−1^·nm^−2^ were applied on all protein Cα atoms for an additional 150 ns. Finally, all production runs were performed without any positional constraints, starting from multiple configurations obtained during equilibration.

Interaction cutoffs were set at 1.2 nm for both Lennard-Jones and Coulombic forces. Long-range electrostatic interactions were calculated using the default reaction-field algorithm. We maintained the system temperature at 323 K using the Nose-Hoover thermostat and controlled pressure at 1 bar, under semi-isotropic conditions with a compressibility of 3 × 10^−4^ bar^−1^, using the Parrinello-Rahman barostat. A longer timestep of 4 fs was set in the PACE simulation.

To explore the relationship between membrane tension and channel activation, we subjected the bilayer XY plane in the PIEZO2 PACE systems to three distinct pressures: +1 bar (corresponding to tensionless bilayer), −2 bar (corresponding to 9.3 ± 0.2 mN m^−1^ membrane tension), and −5 bar (corresponding to 18.0 ± 0.2 mN m^−1^ membrane tension).

### PIEZO2 AA system construction and simulation

The AA PIEZO2 model using CHARMM36m force field^80^ was prepared using the O1 and O2 open state models obtained from PACE simulation. Only the central core domain (containing repeat A, Cap, pore, and C-terminal domain) was kept and embedded in a POPC bilayer and solvated in 0.15M KCl solution. The total AA system contains 622,323 atoms, including 160539 water molecules and 826 POPC lipid molecules. The voltage simulations were conducted using the GROMACS (version 2023.3). The van der Waals interactions were truncated using a cutoff value of 1.2 nm and the force-switching function was applied between the distance range 1.0-1.2 nm. Long-range electrostatics were evaluated using the particle mesh Ewald method and a 1.2 nm cutoff was employed for short range electrostatics. All bonds containing hydrogen atoms were constrained using the LINCS algorithm. A timestep of 2 fs was used for all AA simualtions. The system underwent minimization and equilibration in constant volume and temperature (NVT) and then in constant pressure and temperature (NPT) condition. The Nose-Hoover thermostat and Parrinello-Rahman barostat were employed to maintain 310.15 K and1 atm during the production run. To estimate the conductance of the PIEOZ2 open pores, constant external electric fields of ± 0.6 V were applied in the z-direction. The lateral pressure was kept at −3 bar throughout the simulations.Three replicas of 100 ns simulation was run at each voltage. Ionic current was measured by counting the total number of ions flowing through the pore from the extra cellular to the cytoplasmic side and vice versa.

### Electrophysiology

1µg of plasmids encoding WT mouse PIEZO1 was transfected into 80% confluent HEK293T^PIEZO1KO^ cells in a one well of a 12 well plate as previously described^81^. Cells were re-seeded on 12 mm glass coverslips the next day and used in experiments 36-48 hours after transfection. Patch pipettes were pulled from G150F borosilicate capillaries (Warner Instruments) using a P-97 puller (Sutter Instrument) and heat-polished using a Microforge-MF2 (Narishige). For cell attached pressure clamp experiments shown in Figure 1, patch pipettes were filled with 140 mM KCl, 10 mM HEPES, 10 mM TEA and 2 mM EGTA (pH 7.4). Commercial Hank’s Balanced Saline Solution containing 150 mM NaCl (HBSS, MilliporeSigma, USA) was used as bath solution.

Monovalent cationic saline solutions used in pore size measurements were made of 10 mM HEPES, 10 mM EGTA and 150 mM of a salt solution (NaCl, Methylamine chloride, Dimethylamine chloride, Trimethylamine chloride, or Tetramethylamine chloride). The pH of these saline solutions was adjusted to 7.4 pH using NMDGOH or HCl. Patch pipettes were pulled to a resistance of 2-3 MΩ when filled with these solutions (all chemicals were purchased from Millipore-Sigma). Excised patches were formed in HBSS saline and were sequentially perfused with NaCl, methylamine-Cl, dimethylamine-Cl, trimethylamine-Cl or tetramethylamine-Cl solutions using a pressurized automatic perfusion system (AM-PS8-PR, Sutter Instrument).

The relative permeability of methylated amines (P_MA_) was calculated using a modified Goldman-Hodgkin-Katz (GHK) equation (eq.1) that accounts for the non-zero chloride permeability of PIEZO1 and PIEZO2:

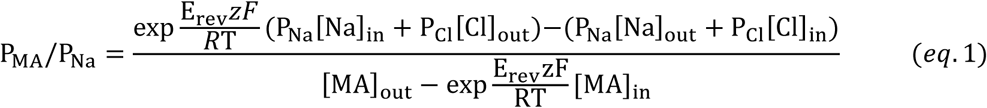

with E_rev_ and *z*, the reversal potential (in Volts) and the valence of the methylamine respectively. *F*, *R* and T are the Faraday constant (96,485 C mol^−1^), the gas constant (8.314 J mol^−1^ K^−1^) and the temperature (in Kelvin), respectively. P_Na_, P_Cl_, are the permeabilities of sodium and chloride ions respectively relative to sodium. [Na]_in_, [Na]_out_, [Cl]_in_, [Cl]_out_, [MA]_in_, and [MA]_out_ are the molar concentrations of indicated saline in intracellular/bath (in) or extracellular/pipette (out) saline, respectively. We have shown that chloride permeability of PIEZO1 and PIEZO2 are voltage dependent^41^. Chloride permeabilities were calculated at each voltage by calculating the ratio of average current recorded in the presence of chloride over sodium ion as sole permeants (P_Cl_ = I_Cl_/I_Na_).

The pore radius was estimated using the following equation based on excluded volume theory^42,47^:

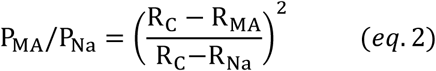

with R_MA_ the radii of methyled amines^82^; R_C_, the minimal radius of the channel pore; and R_Na_, the radius of a sodium ion. Eq. 2 was subjected to a square root transformation, yielding a linear equation used for fitting (OriginPro 2018):

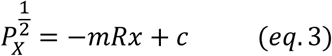

where m = 1/(R_C_-R_Na_) and c = R_C_/(R_C_-R_Na_). According to this equation, the minimum radius of a hypothetical cation with zero permeability corresponds to the pore size. Therefore, the square root of permeability for each tested methylated amine was plotted against its Stokes radius and fitted with a linear function. The x-intercept of this linear fit corresponds to the minimum pore size.

Pressure pulses were delivered through the patch pipette using a high-speed pressure clamp (ALA Scientific). Pressure-induced currents were recorded at 10 kHz in the voltage-clamp mode using an Axopatch 200B amplifier and digitized using a Digidata 1550B (Molecular Devices). Patches were held at +5 mmHg and 0 mV for 20 s between each measurement to allow complete recovery from inactivation^35,83,84^. An Ag/AgCl (3M KCl) reference electrode was used as bath ground. Junction potentials, measured in excised patches from untransfected cells in the current-clamp mode, were below 2 mV at all the tested saline conditions thus were not corrected from I-V curves.

Single channel recordings were obtained as shown earlier^83^ without modification. The number of channels in the membrane patch was estimated by applying a short 10 ms −90 mmHg pressure pulse (Figure 3). The patch was then held at 0 mmHg for 20 s before increasing the pressure to −40 mmHg for 20 s. Single channel data was acquired at 10 KHz and then filtered at 100 Hz to reveal sub conductance levels. All-point histograms and gaussian fitting were done using the Clampfit 10 software (Molecular Devices).

## Data availability

The Gromacs topology files, input and output files for PIEZO2 PACE and AA simulations, data analysis, and final PIEZO2 open state are available to at https://github.com/LynaLuo-Lab/Piezo2/tree/main/piezo2_PACE_simulation. For guidance on the parameters and usage of the PACE force field, please refer to the “PACEm” section available at https://github.com/hanlab-computChem/hanlab/tree/master/PACEm. Electrophysiological data (raw single channel and macroscopic current traces) are available upon request.

## Code availability

The in-house code to generate dome-shaped membrane, are available to at https://github.com/LynaLuo-Lab/Piezo2/tree/main/construct_dome_shaped_membrane.

## Acknowledgements

We thank Ardem Patapoutian (Scripps Research) for the generous gift of HEK293T^PIEZO1KO^ cells and plasmids encoding PIEZO1 and PIEZO2. This work was supported by NIH grants GM130834 (Y.L.L. and J.L.) and National Science Foundation of China Grant No. 21977011 (W.H). The computational resources were provided by the Advanced Cyberinfrastructure Coordination Ecosystem: Services & Support (ACCESS) program, which is supported by National Science Foundation grants #2138259, #2138286, #2138307, #2137603, and #2138296, and the Pittsburgh Supercomputing Center Anton2 allocation MCB170106P, which is supported by NIH Grant GM116961.

## Author Contributions

S.L., A.B., and W.J. performed molecular dynamics simulations. T.W. conducted electrophysiology experiments. Y.L.L, H.W., and J.L. conceived and supervised the project. All authors analyzed the results and prepared the manuscript.

## References

1 Cahalan, S. M. et al. Piezo1 links mechanical forces to red blood cell volume. Elife 4, doi:10.7554/eLife.07370 (2015).

2 Pathak, M. M. et al. Stretch-activated ion channel Piezo1 directs lineage choice in human neural stem cells. Proc. Natl. Acad. Sci. U. S. A. 111, 16148–16153, doi:10.1073/pnas.1409802111 (2014).

3 Gudipaty, S. A. et al. Mechanical stretch triggers rapid epithelial cell division through Piezo1. Nature 543, 118–121, doi:10.1038/nature21407 (2017).

4 Eisenhoffer, G. T. et al. Crowding induces live cell extrusion to maintain homeostatic cell numbers in epithelia. Nature 484, 546–549, doi:10.1038/nature10999 (2012).

5 Zhou, F. et al. Akt/Protein kinase B is required for lymphatic network formation, remodeling, and valve development. Am. J. Pathol. 177, 2124–2133, doi:10.2353/ajpath.2010.091301 (2010).

6 Choi, D. et al. Piezo1 incorporates mechanical force signals into the genetic program that governs lymphatic valve development and maintenance. JCI Insight 4, doi:10.1172/jci.insight.125068 (2019).

7 Li, J. et al. Piezo1 integration of vascular architecture with physiological force. Nature 515, 279–282, doi:10.1038/nature13701 (2014).

8 Ranade, S. S. et al. Piezo1, a mechanically activated ion channel, is required for vascular development in mice. Proc. Natl. Acad. Sci. U. S. A. 111, 10347–10352, doi:10.1073/pnas.1409233111 (2014).

9 Zeng, W. Z. et al. PIEZOs mediate neuronal sensing of blood pressure and the baroreceptor reflex. Science 362, 464–467, doi:10.1126/science.aau6324 (2018).

10 Ranade, S. S. et al. Piezo2 is the major transducer of mechanical forces for touch sensation in mice. Nature 516, 121–125, doi:10.1038/nature13980 (2014).

11 Woo, S. H. et al. Piezo2 is required for Merkel-cell mechanotransduction. Nature 509, 622–626, doi:10.1038/nature13251 (2014).

12 Florez-Paz, D., Bali, K. K., Kuner, R. & Gomis, A. A critical role for Piezo2 channels in the mechanotransduction of mouse proprioceptive neurons. Sci. Rep. 6, 25923, doi:10.1038/srep25923 (2016).

13 Woo, S. H. et al. Piezo2 is the principal mechanotransduction channel for proprioception. Nat. Neurosci. 18, 1756–1762, doi:10.1038/nn.4162 (2015).

14 Wang, L. et al. Structure and mechanogating of the mammalian tactile channel PIEZO2. Nature 573, 225–229, doi:10.1038/s41586-019-1505-8 (2019).

15 Saotome, K. et al. Structure of the mechanically activated ion channel Piezo1. Nature 554, 481–486, doi:10.1038/nature25453 (2018).

16 Guo, Y. R. & MacKinnon, R. Structure-based membrane dome mechanism for Piezo mechanosensitivity. Elife 6, doi:10.7554/eLife.33660 (2017).

17 Zhao, Q. et al. Structure and mechanogating mechanism of the Piezo1 channel. Nature 554, 487–492, doi:10.1038/nature25743 (2018).

18 Zhou, Z. et al. MyoD-family inhibitor proteins act as auxiliary subunits of Piezo channels. Science 381, 799–804, doi:10.1126/science.adh8190 (2023).

19 Haselwandter, C. A. & MacKinnon, R. Piezo’s membrane footprint and its contribution to mechanosensitivity. Elife 7, doi:10.7554/eLife.41968 (2018).

20 Haselwandter, C. A., Guo, Y. R., Fu, Z. & MacKinnon, R. Elastic properties and shape of the Piezo dome underlying its mechanosensory function. Proc. Natl. Acad. Sci. U. S. A. 119, e2208034119, doi:10.1073/pnas.2208034119 (2022).

21 Haselwandter, C. A., Guo, Y. R., Fu, Z. & MacKinnon, R. Quantitative prediction and measurement of Piezo’s membrane footprint. Proc. Natl. Acad. Sci. U. S. A. 119, e2208027119, doi:10.1073/pnas.2208027119 (2022).

22 Jiang, W. et al. Crowding-induced opening of the mechanosensitive Piezo1 channel in silico. Communications Biology 4, 84, doi:10.1038/s42003-020-01600-1 (2021).

23 De Vecchis, D., Beech, D. J. & Kalli, A. C. Molecular dynamics simulations of Piezo1 channel opening by increases in membrane tension. Biophys. J. 120, 1510–1521, doi:10.1016/j.bpj.2021.02.006 (2021).

24 Ozkan, A. D., Wijerathne, T. D., Gettas, T. & Lacroix, J. J. Force-induced motions of the PIEZO1 blade probed with fluorimetry. Cell Rep 42, 112837, doi:10.1016/j.celrep.2023.112837 (2023).

25 Yang, X. et al. Structure deformation and curvature sensing of PIEZO1 in lipid membranes. Nature, doi:10.1038/s41586-022-04574-8 (2022).

26 Mulhall, E. M. et al. Direct observation of the conformational states of PIEZO1. Nature, doi:10.1038/s41586-023-06427-4 (2023).

27 Liu, S. et al. Central pore-opening structure and gating of the mechanosensitive PIEZO1 channel. bioRxiv, 2023.2009.2028.559900, doi:10.1101/2023.09.28.559900 (2023).

28 Murthy, S. E. Deciphering mechanically activated ion channels at the single-channel level in dorsal root ganglion neurons. J. Gen. Physiol. 155, doi:10.1085/jgp.202213099 (2023).

29 Han, W., Wan, C. K., Jiang, F. & Wu, Y. D. PACE Force Field for Protein Simulations. 1. Full Parameterization of Version 1 and Verification. J. Chem. Theory Comput. 6, 3373–3389, doi:10.1021/ct1003127 (2010).

30 Han, W. & Schulten, K. Further Optimization of a Hybrid United-Atom and Coarse-Grained Force Field for Folding Simulations: Improved Backbone Hydration and Interactions between Charged Side Chains. J. Chem. Theory Comput. 8, 4413–4424, doi:10.1021/ct300696c (2012).

31 Li, S., Wu, B. H., Luo, Y. L. & Han, W. Simulations of Functional Motions of Super Large Biomolecules with a Mixed-Resolution Model. J Chem Theory Comput 20, 2228–2245, doi:10.1021/acs.jctc.3c01046 (2024).

32 Verkest, C. et al. Intrinsically disordered intracellular domains control key features of the mechanically-gated ion channel PIEZO2. Nat Commun 13, 1365, doi:10.1038/s41467-022-28974-6 (2022).

33 Lukacs, V. et al. Impaired PIEZO1 function in patients with a novel autosomal recessive congenital lymphatic dysplasia. Nat Commun 6, 8329, doi:10.1038/ncomms9329 (2015).

34 Cox, C. D. et al. Removal of the mechanoprotective influence of the cytoskeleton reveals PIEZO1 is gated by bilayer tension. Nat Commun 7, 10366, doi:10.1038/ncomms10366 (2016).

35 Lewis, A. H. & Grandl, J. Mechanical sensitivity of Piezo1 ion channels can be tuned by cellular membrane tension. Elife 4, doi:10.7554/eLife.12088 (2015).

36 Sukharev, S. I., Sigurdson, W. J., Kung, C. & Sachs, F. Energetic and spatial parameters for gating of the bacterial large conductance mechanosensitive channel, MscL. J. Gen. Physiol. 113, 525–540, doi:10.1085/jgp.113.4.525 (1999).

37 Nomura, T. et al. Differential effects of lipids and lyso-lipids on the mechanosensitivity of the mechanosensitive channels MscL and MscS. Proc. Natl. Acad. Sci. U. S. A. 109, 8770–8775, doi:10.1073/pnas.1200051109 (2012).

38 Fowler, P. W. et al. Membrane stiffness is modified by integral membrane proteins. Soft matter 12, 7792–7803 (2016).

39 Shin, K. C. et al. The Piezo2 ion channel is mechanically activated by low-threshold positive pressure. Sci. Rep. 9, 6446, doi:10.1038/s41598-019-42492-4 (2019).

40 Geng, J. et al. A Plug-and-Latch Mechanism for Gating the Mechanosensitive Piezo Channel. Neuron 106, 438–451 e436, doi:10.1016/j.neuron.2020.02.010 (2020).

41 Wijerathne, T. D., Bhatt, A., Jiang, W., Luo, Y. L. & Lacroix, J. J. Mammalian PIEZO channels rectify anionic currents. Biophys. J., doi:10.1016/j.bpj.2024.11.010 (2024).

42 Dwyer, T. M., Adams, D. J. & Hille, B. The permeability of the endplate channel to organic cations in frog muscle. J Gen Physiol 75, 469–492, doi:10.1085/jgp.75.5.469 (1980).

43 Rychkov, G. Y., Pusch, M., Roberts, M. L., Jentsch, T. J. & Bretag, A. H. Permeation and block of the skeletal muscle chloride channel, ClC-1, by foreign anions. J. Gen. Physiol. 111, 653–665, doi:10.1085/jgp.111.5.653 (1998).

44 Diaz-Franulic, I., Sepulveda, R. V., Navarro-Quezada, N., Gonzalez-Nilo, F. & Naranjo, D. Pore dimensions and the role of occupancy in unitary conductance of Shaker K channels. J. Gen. Physiol. 146, 133–146, doi:10.1085/jgp.201411353 (2015).

45 Voets, T., Janssens, A., Droogmans, G. & Nilius, B. Outer pore architecture of a Ca2+-selective TRP channel. J. Biol. Chem. 279, 15223–15230, doi:10.1074/jbc.M312076200 (2004).

46 Pan, B., Waguespack, J., Schnee, M. E., LeBlanc, C. & Ricci, A. J. Permeation properties of the hair cell mechanotransducer channel provide insight into its molecular structure. J. Neurophysiol. 107, 2408–2420, doi:10.1152/jn.01178.2011 (2012).

47 Chung, M. K., Güler, A. D. & Caterina, M. J. TRPV1 shows dynamic ionic selectivity during agonist stimulation. Nature neuroscience 11, 555–564, doi:10.1038/nn.2102 (2008).

48 Gupta, C., Sarkar, D., Tieleman, D. P. & Singharoy, A. The ugly, bad, and good stories of large-scale biomolecular simulations. Curr. Opin. Struct. Biol. 73, 102338, doi:10.1016/j.sbi.2022.102338 (2022).

49 Liu, S. et al. An intermediate open structure reveals the gating transition of the mechanically activated PIEZO1 channel. Neuron, doi:10.1016/j.neuron.2024.11.020 (2024).

50 Bandarupalli, R., Roth, R., Klipp, R. C., Bankston, J. R. & Li, J. Molecular Insights into Single-Chain Lipid Modulation of Acid-Sensing Ion Channel 3. J Phys Chem B 128, 12685–12697, doi:10.1021/acs.jpcb.4c04289 (2024).

51 Han, Y. et al. Mechanical activation opens a lipid-lined pore in OSCA ion channels. Nature 628, 910–918, doi:10.1038/s41586-024-07256-9 (2024).

52 Reddy, B., Bavi, N., Lu, A., Park, Y. & Perozo, E. Molecular basis of force-from-lipids gating in the mechanosensitive channel MscS. Elife 8, e50486 (2019).

53 Brohawn, S. G., Campbell, E. B. & MacKinnon, R. Physical mechanism for gating and mechanosensitivity of the human TRAAK K+ channel. Nature 516, 126–130 (2014).

54 Jiang, W., Lacroix, J. & Luo, Y. L. Importance of molecular dynamics equilibrium protocol on protein-lipid interaction near channel pore. Biophys Rep (N Y) 2, 100080, doi:10.1016/j.bpr.2022.100080 (2022).

55 Poole, K., Herget, R., Lapatsina, L., Ngo, H. D. & Lewin, G. R. Tuning Piezo ion channels to detect molecular-scale movements relevant for fine touch. Nat Commun 5, 3520, doi:10.1038/ncomms4520 (2014).

56 Qi, Y. et al. Membrane stiffening by STOML3 facilitates mechanosensation in sensory neurons. Nat Commun 6, 8512, doi:10.1038/ncomms9512 (2015).

57 Buyan, A. et al. Piezo1 Forms Specific, Functionally Important Interactions with Phosphoinositides and Cholesterol. Biophys. J., doi:10.1016/j.bpj.2020.07.043 (2020).

58 Ridone, P. et al. Disruption of membrane cholesterol organization impairs the activity of PIEZO1 channel clusters. J. Gen. Physiol. 152, doi:10.1085/jgp.201912515 (2020).

59 Romero, L. O. et al. Dietary fatty acids fine-tune Piezo1 mechanical response. Nat Commun 10, 1200, doi:10.1038/s41467-019-09055-7 (2019).

60 Romero, L. O. et al. A dietary fatty acid counteracts neuronal mechanical sensitization. Nat Commun 11, 2997, doi:10.1038/s41467-020-16816-2 (2020).

61 Gabrielle, M., Yudin, Y., Wang, Y., Su, X. & Rohacs, T. Phosphatidic acid is an endogenous negative regulator of PIEZO2 channels and mechanical sensitivity. Nat Commun 15, 7020, doi:10.1038/s41467-024-51181-4 (2024).

62 Borbiro, I., Badheka, D. & Rohacs, T. Activation of TRPV1 channels inhibits mechanosensitive Piezo channel activity by depleting membrane phosphoinositides. Sci Signal 8, ra15, doi:10.1126/scisignal.2005667 (2015).

63 Narayanan, P. et al. Myotubularin related protein-2 and its phospholipid substrate PIP2 control Piezo2-mediated mechanotransduction in peripheral sensory neurons. Elife 7, doi:10.7554/eLife.32346 (2018).

64 Shi, J. et al. Sphingomyelinase Disables Inactivation in Endogenous PIEZO1 Channels. Cell Rep 33, 108225, doi:10.1016/j.celrep.2020.108225 (2020).

65 Tsuchiya, M. et al. Cell surface flip-flop of phosphatidylserine is critical for PIEZO1-mediated myotube formation. Nat Commun 9, 2049, doi:10.1038/s41467-018-04436-w (2018).

66 Shan, Y. G., X.; Zhang, M.; Chen, M; Li, Y.; Zhang, M.; Pei, D. Structure of human PIEZO1 and its slow inactivating channelopathy mutant. BioRxiv, doi:10.1101/2024.07.14.603468 (2024).

67 Szczot, M. et al. PIEZO2 mediates injury-induced tactile pain in mice and humans. Sci. Transl. Med. 10, doi:10.1126/scitranslmed.aat9892 (2018).

68 Murthy, S. E. et al. The mechanosensitive ion channel Piezo2 mediates sensitivity to mechanical pain in mice. Sci. Transl. Med. 10, doi:10.1126/scitranslmed.aat9897 (2018).

69 Dubin, A. E. et al. Inflammatory signals enhance piezo2-mediated mechanosensitive currents. Cell Rep 2, 511–517, doi:10.1016/j.celrep.2012.07.014 (2012).

70 Servin-Vences, M. R. et al. PIEZO2 in somatosensory neurons controls gastrointestinal transit. Cell 186, 3386–3399 e3315, doi:10.1016/j.cell.2023.07.006 (2023).

71 Marshall, K. L. et al. PIEZO2 in sensory neurons and urothelial cells coordinates urination. Nature, doi:10.1038/s41586-020-2830-7 (2020).

72 Jiang, W. et al. Structural and thermodynamic framework for PIEZO1 modulation by small molecules. Proc. Natl. Acad. Sci. U. S. A. 120, e2310933120, doi:10.1073/pnas.2310933120 (2023).

73 Botello-Smith, W. M. et al. A mechanism for the activation of the mechanosensitive Piezo1 channel by the small molecule Yoda1. Nature Communications 10, 4503, doi:10.1038/s41467-019-12501-1 (2019).

74 Lacroix, J. J., Botello-Smith, W. M. & Luo, Y. Probing the gating mechanism of the mechanosensitive channel Piezo1 with the small molecule Yoda1. Nature Communications 9, 2029, doi:10.1038/s41467-018-04405-3 (2018).

75 Webb, B. & Sali, A. Comparative Protein Structure Modeling Using MODELLER. Curr Protoc Protein Sci 86, 2 9 1–2 9 37, doi:10.1002/cpps.20 (2016).

76 Eastman, P. et al. OpenMM 7: Rapid development of high performance algorithms for molecular dynamics. PLoS Comput Biol 13, e1005659, doi:10.1371/journal.pcbi.1005659 (2017).

77 Periole, X., Cavalli, M., Marrink, S. J. & Ceruso, M. A. Combining an Elastic Network With a Coarse-Grained Molecular Force Field: Structure, Dynamics, and Intermolecular Recognition. J. Chem. Theory Comput. 5, 2531–2543, doi:10.1021/ct9002114 (2009).

78 Hess, B., Kutzner, C., van der Spoel, D. & Lindahl, E. GROMACS 4: Algorithms for Highly Efficient, Load-Balanced, and Scalable Molecular Simulation. J Chem Theory Comput 4, 435–447, doi:10.1021/ct700301q (2008).

79 Mark James Abraham, T. M., Roland Schulz, Szilárd Páll, Jeremy C. Smith, Berk Hess, Erik Lindahl,. GROMACS: High performance molecular simulations through multi-level parallelism from laptops to supercomputers,. SoftwareX, 1–2,, 19–25,, doi:10.1016/j.softx.2015.06.001 (2015).

80 Huang, J. et al. CHARMM36m: an improved force field for folded and intrinsically disordered proteins. Nat. Methods 14, 71–73, doi:10.1038/nmeth.4067 (2017).

81 Wijerathne, T. & Lacroix, J. Voltage-clamp fluorometry to record flow-activated PIEZO1 currents and fluorometric signals. STAR Protoc 5, 102789, doi:10.1016/j.xpro.2023.102789 (2024).

82 Liu, D. M. & Adams, D. J. Ionic selectivity of native ATP-activated (P2X) receptor channels in dissociated neurones from rat parasympathetic ganglia. J. Physiol. 534, 423–435, doi:10.1111/j.1469-7793.2001.00423.x (2001).

83 Wijerathne, T. D., Ozkan, A. D. & Lacroix, J. J. Yoda1’s energetic footprint on Piezo1 channels and its modulation by voltage and temperature. Proc. Natl. Acad. Sci. U. S. A., e2202269119, doi:doi.org/10.1073/pnas.2202269119 (2022).

84 Wu, J. et al. Inactivation of Mechanically Activated Piezo1 Ion Channels Is Determined by the C-Terminal Extracellular Domain and the Inner Pore Helix. Cell Rep 21, 2357–2366, doi:10.1016/j.celrep.2017.10.120 (2017).

