## Supplementary Information for "A two-step clockwork mechanism opens a proteo-lipidic pore in PIEZO2 channels"

### Table S1. All-atom MD simulated KCl conductance for O1 and O2 states.

| State | Voltage (V) | Replica | Time (ns) | K+ | K+ Current (pA) | Cl- | Cl- Current (pA) | Total Current (pA) | Conductance (pS) |
|---|---|---|---|---|---|---|---|---|---|
| **O1** | -0.6 | R-1 | 100 | 2 | -3.2 | 1 | -1.6 | -4.8 | 8 |
| | -0.6 | R-2 | 100 | 1 | -1.6 | 1 | -1.6 | -3.2 | 5.3 |
| | -0.6 | R-3 | 100 | 3 | -4.8 | 2 | -3.2 | -8.0 | 13.3 |
| | | | | | | | | | **8.9 ± 4.1** |
| | 0.6 | R-1 | 100 | 3 | 4.8 | 0 | 0.0 | 4.8 | 8.0 |
| | 0.6 | R-2 | 100 | 4 | 6.4 | 3 | 4.8 | 11.2 | 18.7 |
| | 0.6 | R-3 | 100 | 2 | 3.2 | 1 | 1.6 | 4.8 | 8.0 |
| | | | | | | | | | **11.6 ± 6.2** |
| **O2** | -0.6 | R-1 | 100 | 12 | -19.2 | 1 | -1.6 | -20.8 | 34.7 |
| | -0.6 | R-2 | 100 | 13 | -20.8 | 1 | -1.6 | -22.4 | 37.3 |
| | -0.6 | R-3 | 100 | 11 | -17.6 | 1 | -1.6 | -19.2 | 32.0 |
| | | | | | | | | | **34.7 ± 2.7** |
| | 0.6 | R-1 | 100 | 9 | 14.4 | 7 | 11.2 | 25.6 | 42.7 |
| | 0.6 | R-2 | 100 | 6 | 9.6 | 5 | 8.0 | 17.6 | 29.3 |
| | 0.6 | R-3 | 100 | 10 | 16.0 | 6 | 9.6 | 25.6 | 42.7 |
| | | | | | | | | | **38.2 ± 7.7** |

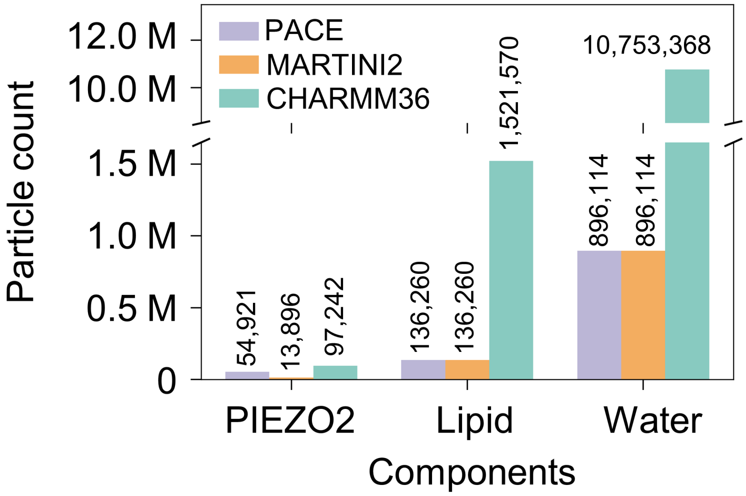


**Fig.S1a Number of particles in three PIEOZ2 models.**

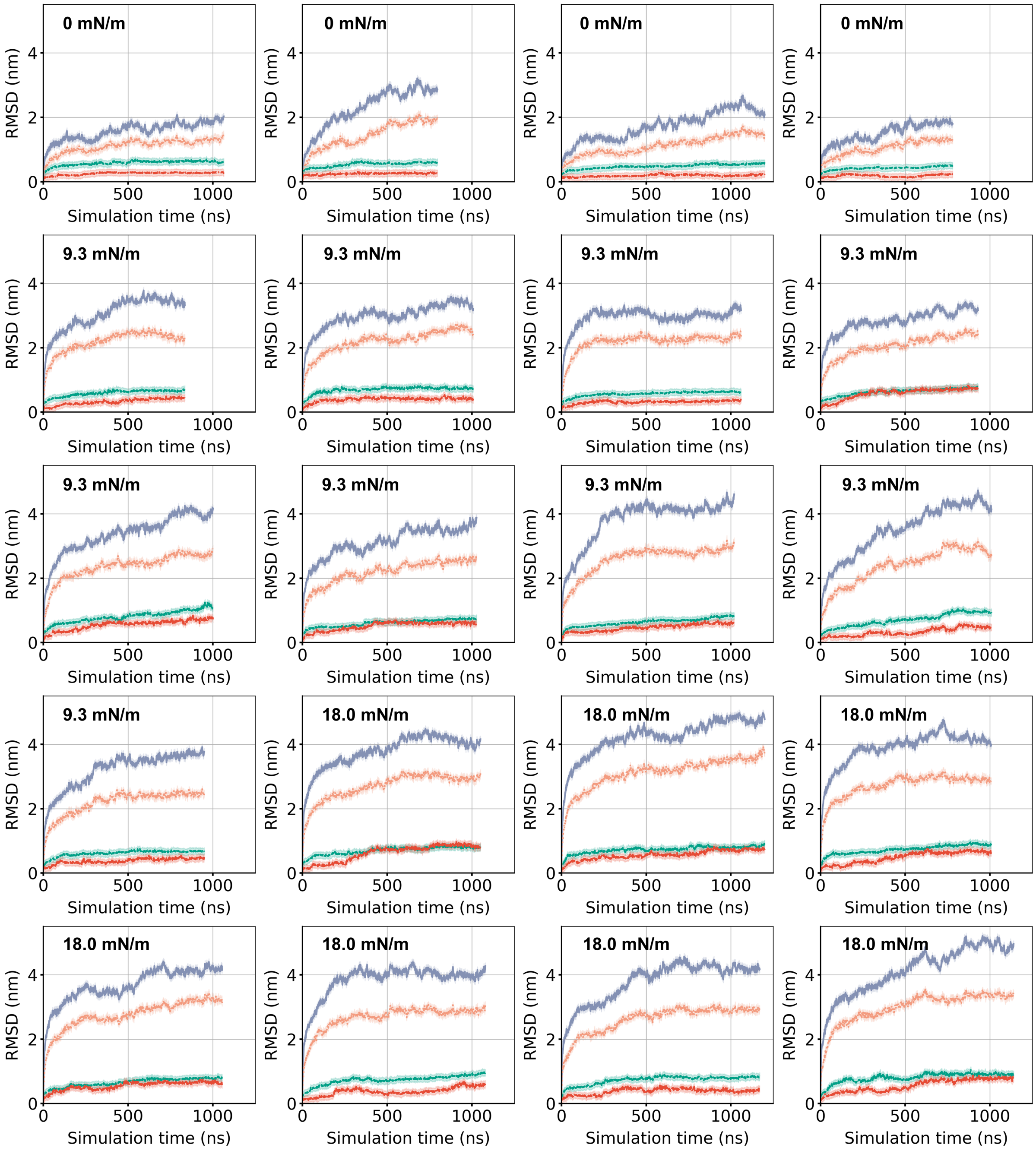


**Fig.S1b RMSD analysis of PIEZO2 global structure to key internal domains.** RMSD values for the protein Cα atoms was calculated from 4 replicas at 0 mN/m, 9 replicas at 9.3 mN/m, and 7 replicas at 18.0 mN/m. The navy blue represents the global PIEZO2 structure, orange represents to a portion of PIEZO2 (excluding the repeat G, H, and I regions), green represents the pore domain, and red represents the inner pore helix (TM38).

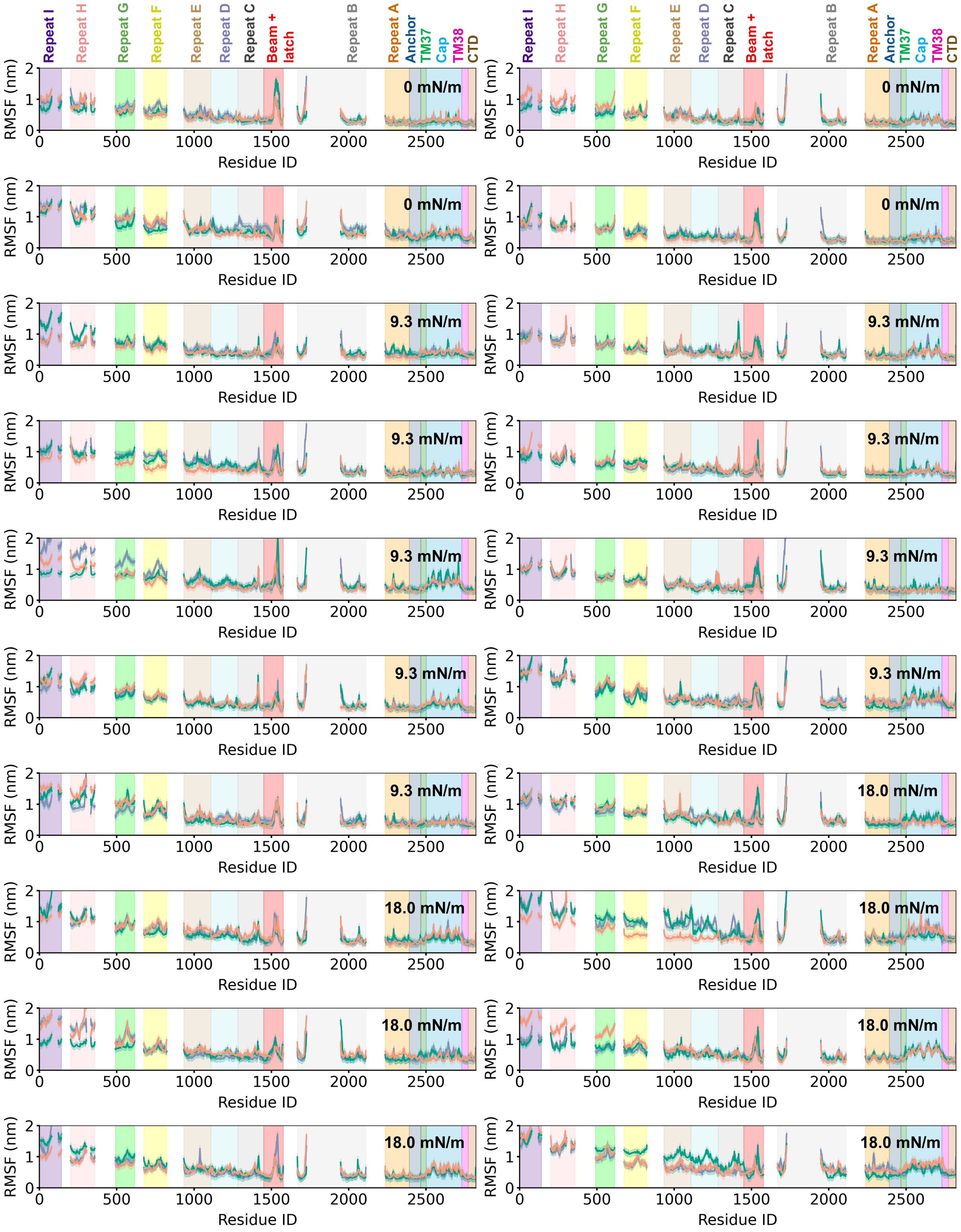


**Fig.S2 Average RMSF analysis of PIEZO2.** The average RMSF of PIEZO2 for each simulation was calculated relative to the conformation at 500 ns in that specific simulation. The three chains of PIEZO2 were represented in navy blue, orange, and green, respectively. Only the conformations after 500 ns of each MD trajectory were included in this analysis.

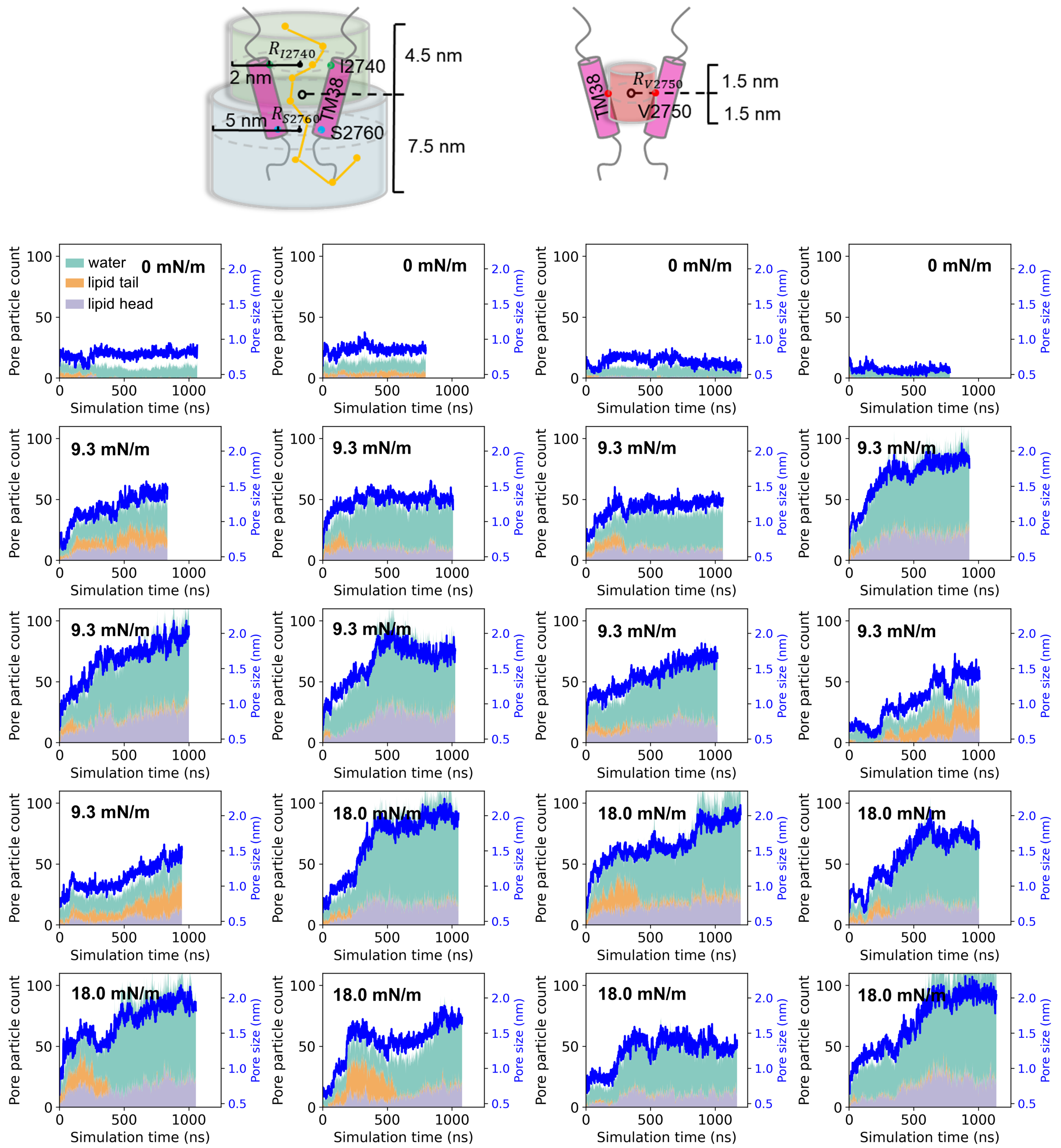


**Fig. S3a. Pore particle count and pore size.** Pore particle count under different membrane tensions. Light lavender gray, light orange, and light mint green represent lipid head particles, lipid tail particles, and water particles, respectively. The blue line indicates the pore size. **Top left:** illustration of the calculation of Martini water pathways in the pore region. The pathways of water particles (yellow spheres) within the pore were determined by analyzing their positions in predefined cylindrical regions. The light green cylinder represents the N-terminal region of TM38, with its radius calculated as the average distance between the Cα atoms of the three I2740 residues and their centroid, plus 2 nm. Similarly, the light blue cylinder represents the C-terminal region of TM38, with its radius calculated as the average distance between the Cα atoms of the three S2760 residues and their centroid, plus 5 nm. Along the Z-axis, the water particles were restricted to a region spanning 4.5 nm above to 7.5 nm below the centroid of the three TM38 helices. For each simulation frame (sampled every 1 ns), the coordinates and IDs of water particles residing within the light green and light blue cylindrical regions were recorded. A water pathway was defined as the continuous trajectory of a water particle appearing in consecutive frames as it moved between the light green and light blue cylindrical regions or vice versa. **Top right:** illustration of pore particle count calculation. The number of particles in the pore was calculated within a defined cylindrical region. The radius of the cylinder was determined as the average distance between the Cα particles of three Val2750 residues and their centroid, while the height extended 1.5 nm above and below the centroid along the Z-axis.

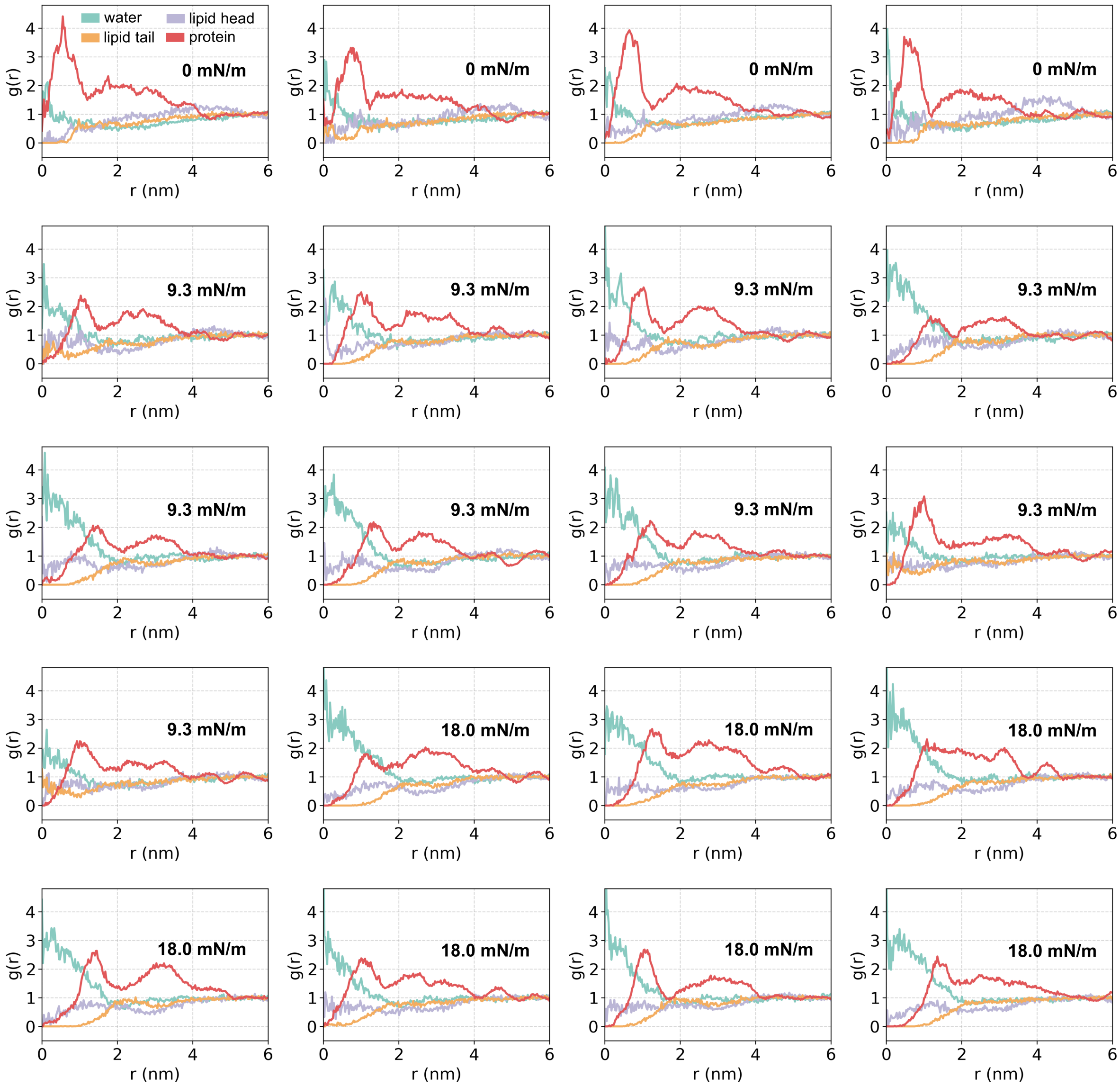


**Fig.S3b. Radial distribution function (RDF) of lipid head (light lavender gray), lipid tail (light orange), water (light mint) and protein (red) around pore center for different membrane tension systems.** The pore center is defined as the center of geometry (COG) of three V2750 Cα particles. The RDF is computed using the pore center as the cylindrical center. The cylinder has a radius of 6 nm and heights of 7 nm (lipid), 5 nm (water), and 3 nm (protein), respectively. For each target particle, the radial distance to the cylinder axis (i.e., the perpendicular distance from the particle to the vertical cylinder axis) is calculated. The RDF, $g(r)$ is defined as $g(r) = \frac{\Delta N(r)}{\rho_{ref} A(r)}$, where $\Delta N(r)$: the number of particles within the annular shell between $r$ and $r + \Delta r$; $A(r)$: the area of the annular shell, calculated as $A(r) = \pi[(r + \Delta r)^2 - r^2]$; $\rho_{ref}$: the average density of particles in the reference range (4-6 nm). Only the conformations after 500 ns of each MD trajectory were included in the statistical analysis.

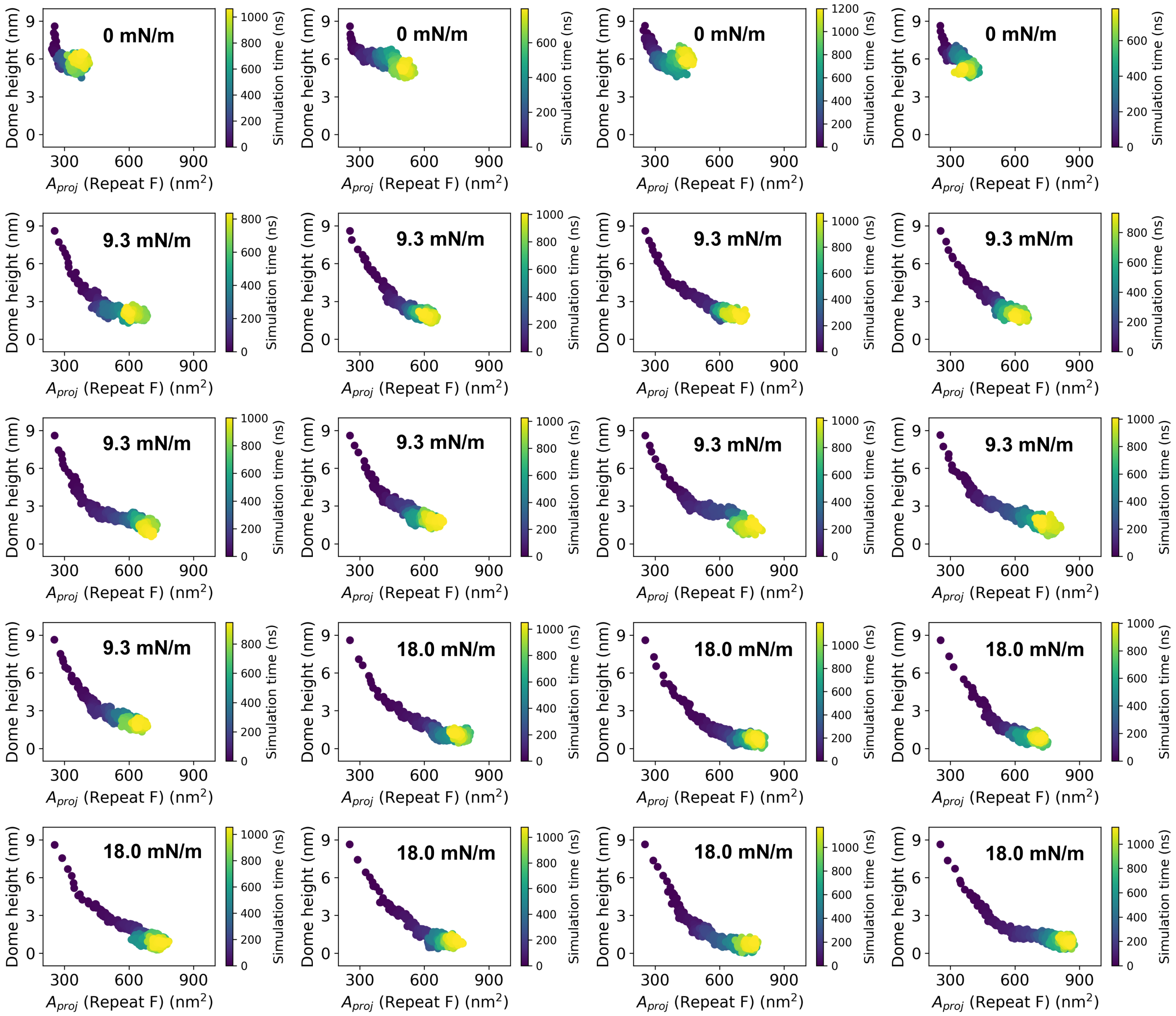


**Fig.S3c. Time evolution of dome height vs. projected area.**

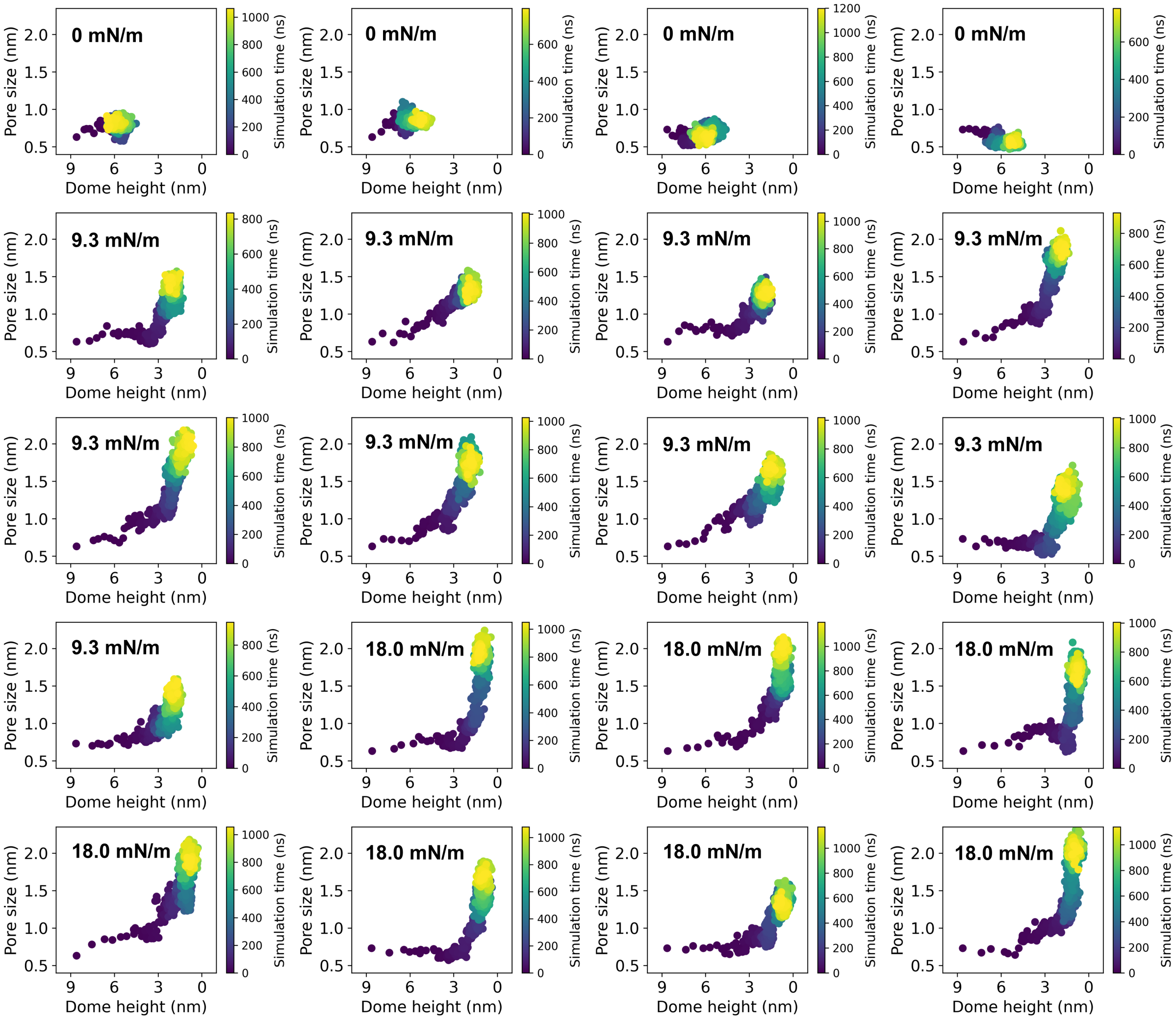


**Fig.S3d. Time evolution of pore size vs. dome height.**

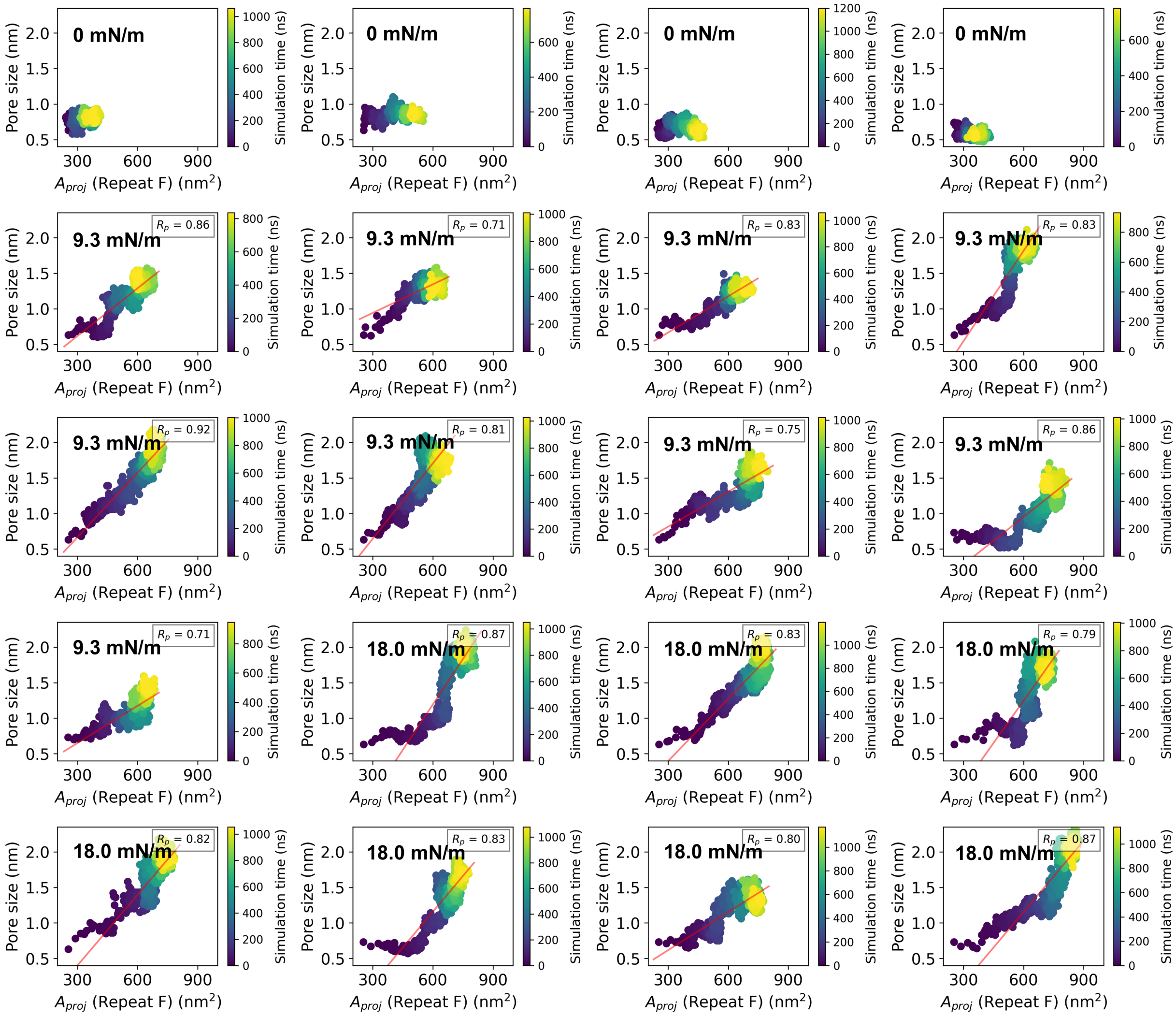


**Fig.S3e. Time evolution of pore size vs. projected area.**

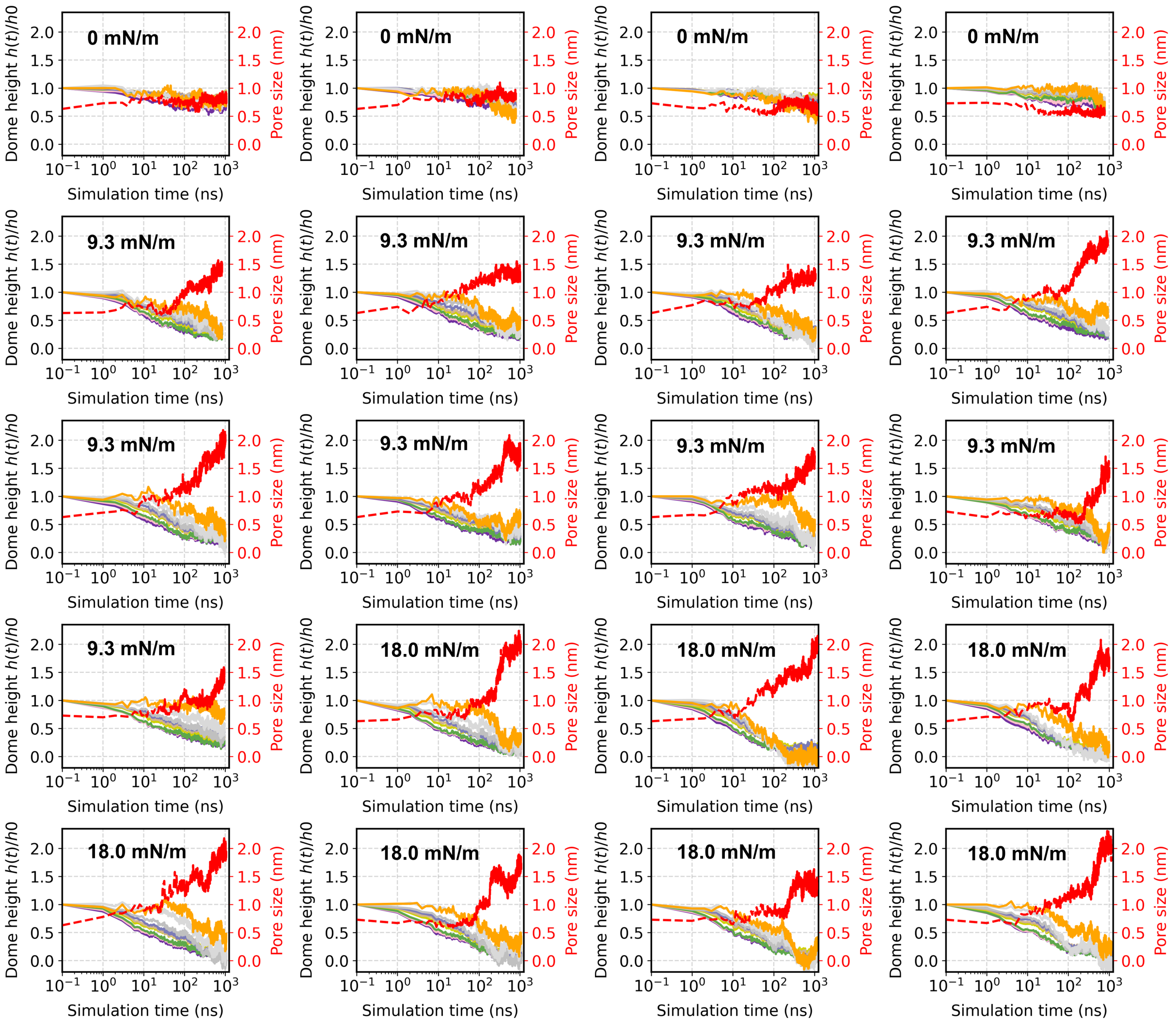


**Fig.S3f. Time evolution of pore size vs. dome height for each transmembrane helical repeat.** Repeat I, H, G, F, E, D, C, B, and A are represented by solid lines in the following colors: purple, pink, green, yellow, tan, blue, silver, light gray, and orange, respectively. The pore size is represented by a red dashed line.

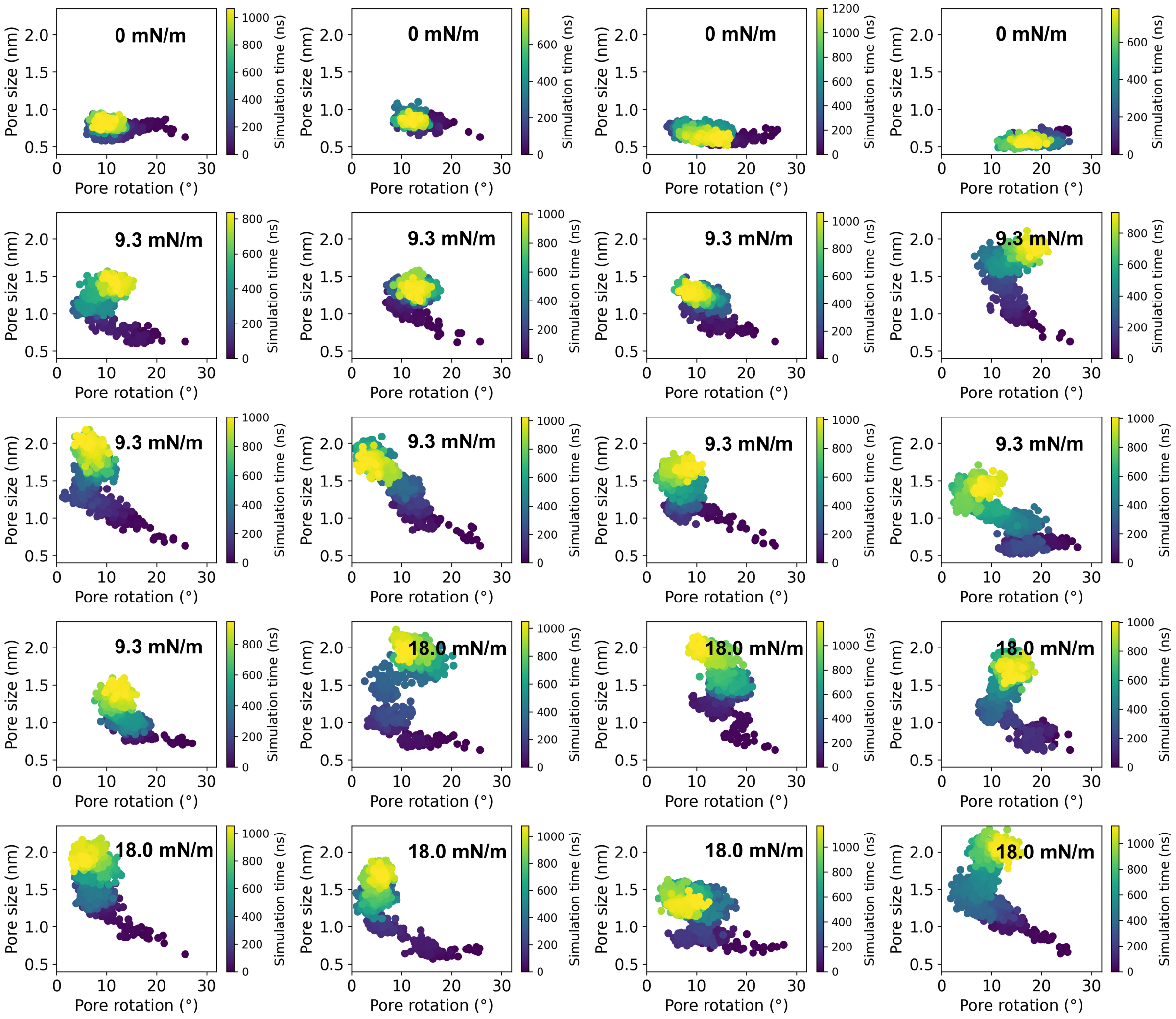


**Fig.S4a. Time evolution of pore size vs. pore rotation.**

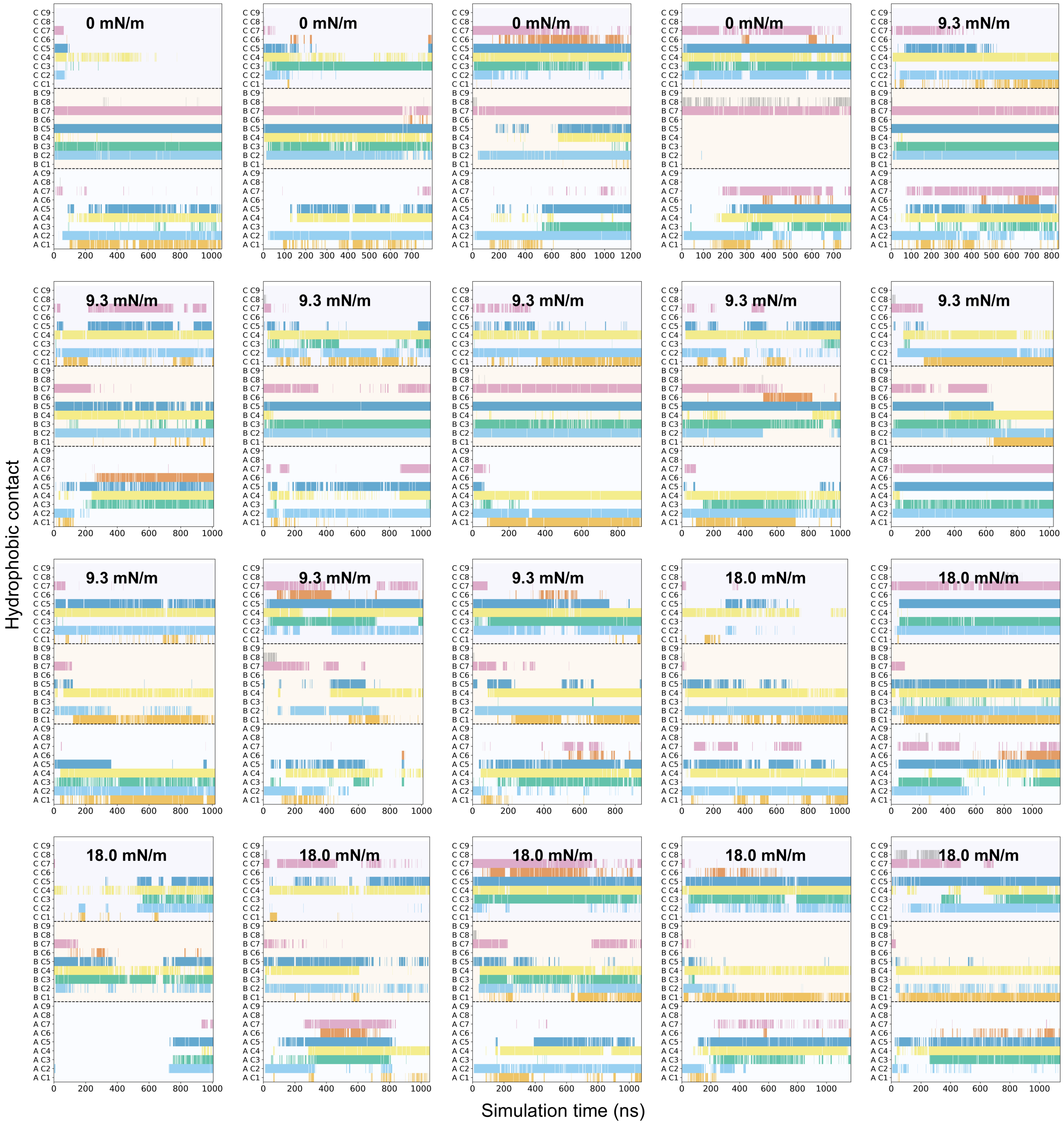


**Fig.S4b. Hydrophobic contact between TM37 and TM38.** A contact is defined when the distance between any pair of particles from two residues is less than 0.45 nm. The light blue, light yellow, and light purple background shading indicates the three chains of PIEZO2. Contact pairs are color-coded as follows: C1 (M2741-F2488) in orange, C2 (M2741-F2492) in sky blue, C3 (M2741-M2493) in green, C4 (F2744-F2488) in yellow, C5 (F2744-F2492) in dark blue, C6 (F2744-M2493) in red, C7 (F2748-F2488) in pink, C8 (F2748-F2492) in gray, and C9 (F2748-M2493) in black. A pair is displayed in its corresponding color if a contact is present.

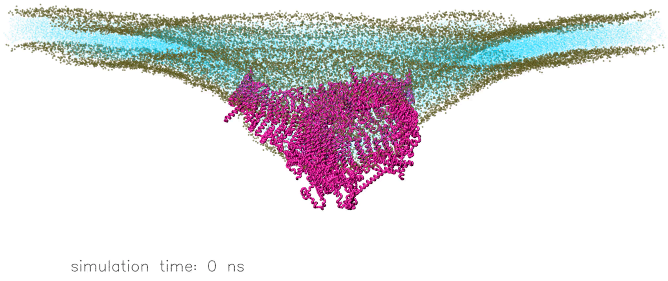


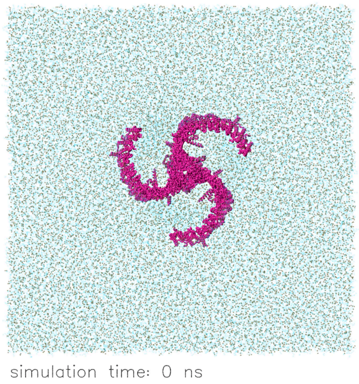


**Movie 1. Molecular dynamics simulation of the full-length PIEZO2 channel embedded in a POPC membrane patch of ~3900 nm$^2$ under 9.3 mN/m membrane tension.** Color code in side view (left) and top view (right): Protein in magenta, lipid in cyan except phosphate group in tan.

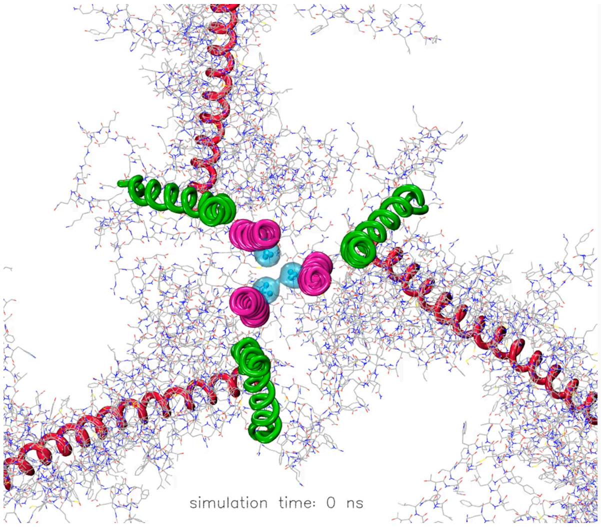


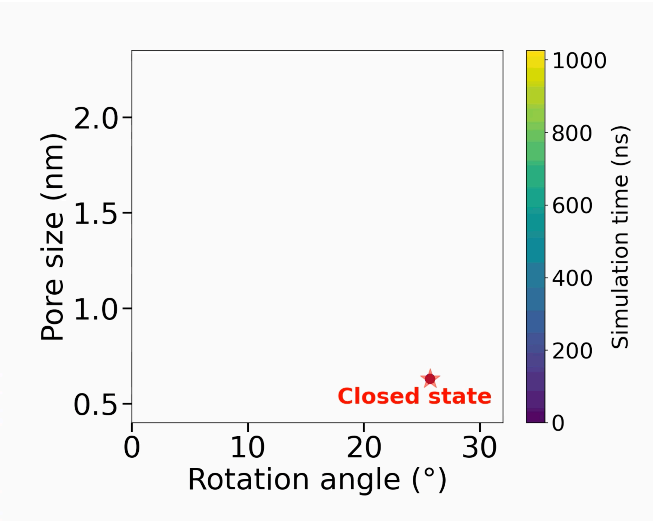


**Movie 2. Clockwork gating motions for PIEZO2 opening at 9.3 mN/m. Left:** Top view of the pore helix rotation. The protein backbone of the inner pore helices (TM38) are shown in magenta, the outer pore helices (TM37) are shown in green, and the beam helices are shown in red. V2750 sidechains are in blue. The rest of the proteins are shown in line mode with atom color code: red oxygen, blue nitrogen, and grey carbon. **Right:** The time evolution of the relative angle between the inner and outer triangles, respectively formed by the trios of TM38 and TM37 helices, are plotted against the pore size at 9.3 mN m$^{-1}$ tension.
